# Multiple targets of balancing selection in *Leishmania donovani* complex parasites

**DOI:** 10.1101/2021.03.02.433528

**Authors:** Cooper Alastair Grace, Sarah Forrester, Vladimir Costa Silva, Aleksander Aare, Hannah Kilford, Yen Peng Chew, Sally James, Dorcas L. Costa, Jeremy C. Mottram, Carlos C. H. N. Costa, Daniel C. Jeffares

## Abstract

The *Leishmania donovani* species complex are the causative agents of visceral leishmaniasis, which cause 20-40,000 fatalities a year. Here, we conduct a screen for balancing selection in this species complex. We used 387 publicly-available *L. donovani* and *L. infantum* genomes, and sequence 93 isolates of *L. infantum* from Brazil to describe the global diversity of this species complex. We identify five genetically-distinct populations that are sufficiently represented by genomic data to search for signatures of selection. We find that signals of balancing selection are generally not shared between populations, consistent with transient adaptive events, rather than long-term balancing selection. We then apply multiple diversity metrics to identify candidate genes with robust signatures of balancing selection, identifying a curated set of 19 genes with robust signatures. These include zeta toxin, nodulin-like and flagellum attachment proteins. This study highlights the extent of genetic divergence between *L. donovani complex* parasites and provides genes for further study.

## Introduction

Intracellular Leishmania parasites cause the neglected infectious disease leishmaniasis in over 80 countries. Visceral leishmaniasis (VL) is the most severe form of the disease, caused by *Leishmania donovani* and *Leishmania infantum*. Yearly cases of VL are estimated at a minimum of 50,000, with a fatality of ≥ 95% if untreated, and occur primarily in the Indian subcontinent (ISC), Bangladesh, Sudan, South Sudan, Ethiopia and Brazil (World Health Organisation 2020). After transmission by sand flies, *Leishmania* promastigotes are taken up by macrophages, and develop into amastigotes which proliferate. These processes require specific adaptations to different environments, such as evasion and active modulation of mammalian host or sand fly vector immune responses (Atayde et al. 2016; Dong et al. 2019). *Leishmania* species contain genomes that are primarily diploid and sexually-recombining. Amongst their unusual features are the use of constitutively-transcribed polycistronic genes and supernumerary chromosomes with unstable polidy (Dumetz et al. 2017).

In *Plasmodium* parasites that have been studied more intensively, some host-interacting genes maintain high genetic diversity due to balancing selection (BS) within populations (Ochola-Oyier et al. 2019), in some cases with a bias towards polymorphic residues to be exposed on surface epitopes (Guy et al. 2018). Given the competitive interaction between *Leishmania* cells and host immune cells, balancing selection may also be present in this parasite. Other drivers of balancing selection, such as heterozygote advantage (overdominance) or alleles that confer fitness differentially in the sand fly vector and the mammal host are also possible. These processes are expected to generate similar genetic signals (Charlesworth 2006). In all these scenarios, genomic signatures of BS can highlight genes that are important for transmission, host immune evasion or ecological adaptation.

Thus far, there have been no published studies of BS in *Leishmania* species. Here, we use genome data from 477 clinical isolates from the *L. donovani* species complex (*L. infantum* or *L. donovani*) from East Africa, the Indian subcontinent (ISC) and Brazil to identify five populations that are well-represented by genome data. Using a variety of metrics, we search for signals of balancing selection within these populations. We identify multiple strong signatures of balancing selection. Signatures are generally unique to a single population consistent with adaptive divergence between populations.

## Results

### *L. donovani* complex genome data and population structure

In this study we used population-scale genomic data from *L. donovani* species complex covering the main global foci of East Africa, the Indian subcontinent (ISC), Brazil and Europe. We utilised 229 *L. donovani* isolates from the ISC (Imamura et al. 2016), 43 *L. donovani* isolates from Ethiopia (Zackay et al. 2018), 25 *L. infantum* isolates from Brazil (Carnielli et al. 2018), and 87 *L. donovani* isolates from a variety of locations including Sudan (14 strains), France (6) and Israel (10) (Franssen et al. 2020). Additionally, we sequenced 93 *L. infantum* isolates from Piauí state, Brazil (Figure 1, **Supplementary Table 1**). This produced a data set of 477 sequenced isolates from the *L. donovani* complex. To detect genetic variants in these genomes we mapped reads from all isolates to the *L. donovani* BPK282A1 reference genome, and applied variant calling methods and filtering to identify single-nucleotide polymorphisms (SNPs) and insertion/deletion polymorphisms (indels). In this data set we identify 339,367 SNPs and 14,383 indels.

**Figure 1:**
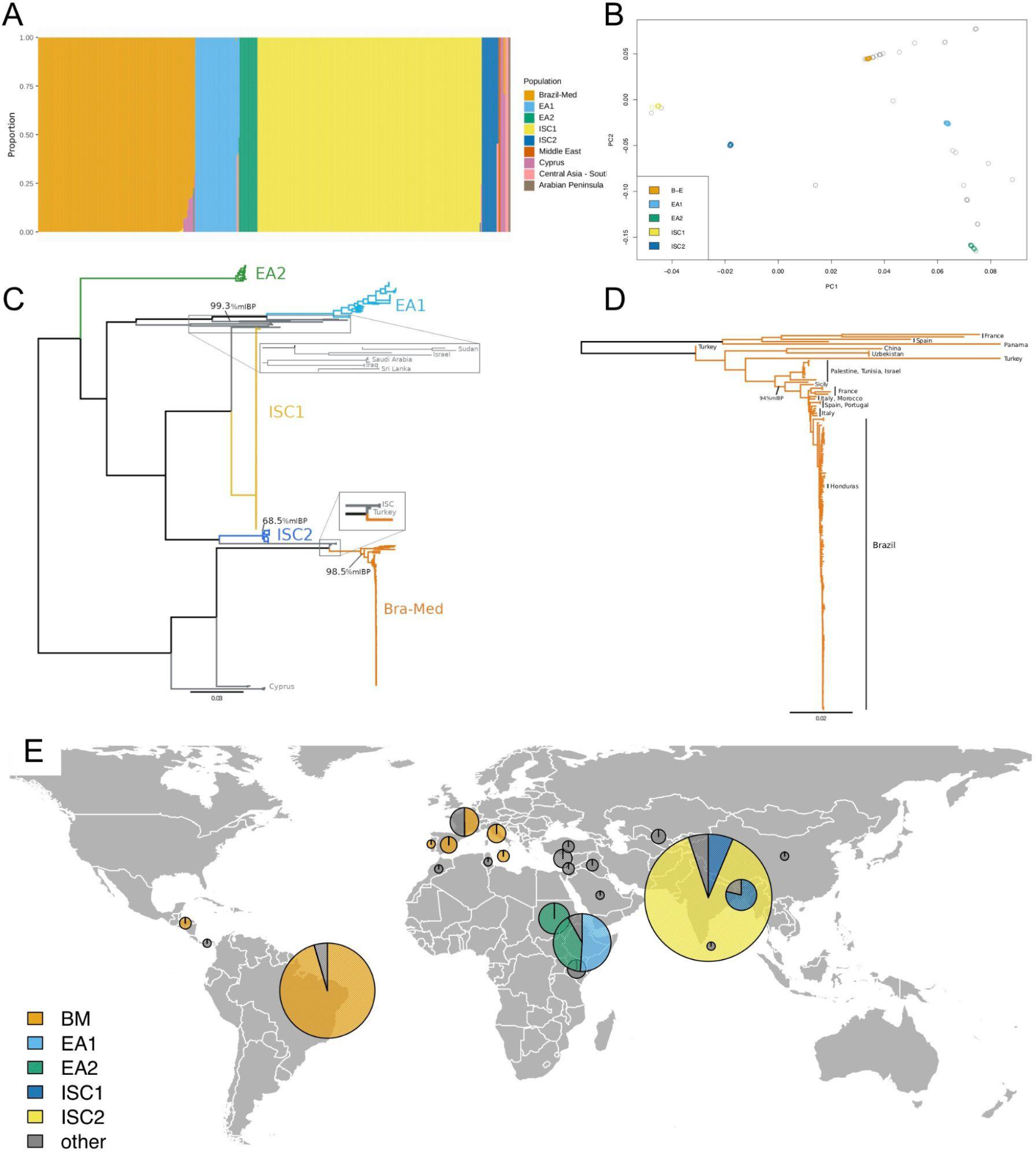
Population structure of the *L. donovani* complex. **Panel A:** ADMIXTURE analysis indicated between 8-11 populations, here *K*=9. Cross validation error values are available as Supplementary Figure 1. ADMIXTURE plots for *K*=8, 10 and 11 populations available as Supplementary Figure 2. **Panel B**: Principal component analysis (PCA). Strains are coloured as for part A. Isolates in grey were not confidently assigned to one of the five major populations (Brazil-Med, EA1, EA2, ISC1 and ISC2) by ADMIXTURE. **Panel C**: Maximum likelihood phylogeny, based upon a SNP alignment of 477 sequences with 283,378 variable sites. All visible branches are maximally supported (100% mlBP) unless indicated. The tree is midpoint rooted, with the unrooted tree available as Supplementary Figure 3. The scale bar represents the number of nucleotide changes per site. Country names in grey indicate origins of the minor populations. **Panel D**: Brazil-Mediterranean tree of *L. infantum* strains, based upon a SNP alignment of 158 sequences with 81,018 variable sites, midpoint rooted. Scale bar and support are as in Panel C. A single sample isolated in China (Franssen et al. 2020), is the only demographic exception in the Brazil-Mediterranean sample collection, and is indicative of movement of parasites. Data and tree files are available in supplementary documents. **Panel E:** Locations of samples used in this study. Pie charts show the number of samples from each location that are confidently assigned to one of the five major populations, with a radius proportional to the number of samples from each location.

We used the ADMIXTURE clustering tool (Alexander and Lange 2011) to assign isolates to populations. This analysis indicated that this collection can be clustered into between 8 and 11 populations (Figure 1, and Supplementary Figures 1-2). The majority of these isolates could be assigned with ≥ 99% confidence to one of five relatively well-sampled populations (Figure 1). Principal component and phylogenetic analysis showed consistent results. These five populations included two from the Indian Sub-Continent (ISC1, ISC2), two from East Africa (EA2 from North Ethiopia/Sudan, and EA1 which corresponds to a population from South Ethiopia/Kenya; Gelanew 2010) and a Brazil-Mediterranean population (BM). The remaining isolates were assigned to populations of <6 isolates (*n*=44).

These five populations have largely independent ancestries. Fixation index (F_ST_) values range from 0.27 to 0.90 (**Supplementary Table 2**). As has been observed previously (Gelanew 2010), the two East African populations and the older ISC population (ISC2) are approximately equidistant, with F_ST_ in the range of ∼0.3. Larger F_ST_ values appear to be due to genetic divergence from the newly-emerged ISC1 and Brazil-Mediterannean populations. Only 7% of polymorphic sites are shared between two or more populations. We note that rare hybrids have been shown to occur between *L. donovani* complex populations in both East Africa and Turkey (Rogers et al. 2014; Cotton et al. 2020). We do not include *L. donovani* complex hybrids from Turkey (Rogers et al. 2014) in our analysis, because hybrid populations may contain balanced alleles that give an appearance of BS. This, and the under-sampling of VL-endemic regions between Europe and India, render our data unsuited to studying the true extent of global gene flow in this species, so we do not analyse this further here.

Phylogenetic analysis provides some qualitative insight to the history of these species (Figure 1, **Panels C and D**). The basal position of EA2 and early-branching position of EA1 support the relative age of these populations in East Africa, as does the high genetic diversity in this region and genetic distance between these populations, consistent with previous studies (Gelanew et al. 2010; Ferreira et al. 2012; Gelanew et al. 2014; Teixeira et al. 2017; Zackay et al. 2018; Cotton et al. 2020; Franssen et al. 2020). The high nucleotide diversity of EA1 (Table 1) is reflected in the branch lengths in this clade of the phylogeny. In contrast, the smaller Indian subcontinent population, identified as ISC1 here (equivalent to the ISC5 group identified by (Imamura et al. 2016), produces short terminal branches in the phylogeny and lower genetic diversity (Table 1, Figure 2), consistent with previous genomic analyses indicating that is an emergent population (Imamura et al. 2016). Epidemiological evidence indicates that this population arose in the 1970s after the malaria elimination programme (Dye and Wolpert 1988; Bhattacharya et al. 2006; Thakur 2007; Muniaraj 2014; Dhiman and Yadav 2016). The 93 Brazilian isolates we examined, which are mostly from Piauí state in north west Brazil, cluster within isolates originating from Mediterranean countries (Figure 1, **Panel D**), consistent with a relatively recent European origin for *L. infantum* in Brazil (<400 years; Kuhls et al. 2011). Short branches in the Brazilian clade (Figure 1, **Panel C-D**), low genetic diversity and an abundance of rare alleles (Figure 2) are all consistent with a relatively recent population bottleneck and an expanding population. In contrast, both East African and populations and the older population from the Indian subcontinent (ISC2) show higher genetic diversity, and are likely to have been maintained as larger populations for longer periods of time.

**Table 1.** Population statistics for *L. donovani* complex populations.

| Pop | Source | Isolates | No. of<br>'pure' | SNPs | Indels | $\pi$<br>$\times 10^{-6}$ | Tajima<br>*D* | Mean<br>MAF |
| --- | --- | --- | --- | --- | --- | --- | --- | --- |
| EA1 | East<br>Africa | 41 | 41 | 3033 | 705 | 424 | 0.70 | 0.23 |
| EA2 | East<br>Africa | 18 | 18 | 970 | 575 | 87 | -0.20 | 0.14 |
| ISC1 | India | 225 | 211 | 2689 | 551 | 84 | -0.23 | 0.17 |
| ISC2 | India | 15 | 15 | 3103 | 716 | 6.3 | -1.04 | 0.009 |
| BM | Brazil,<br>Med. | 133 | 127 | 8886 | 1578 | 28 | -1.42 | 0.02 |

**Figure 2.**
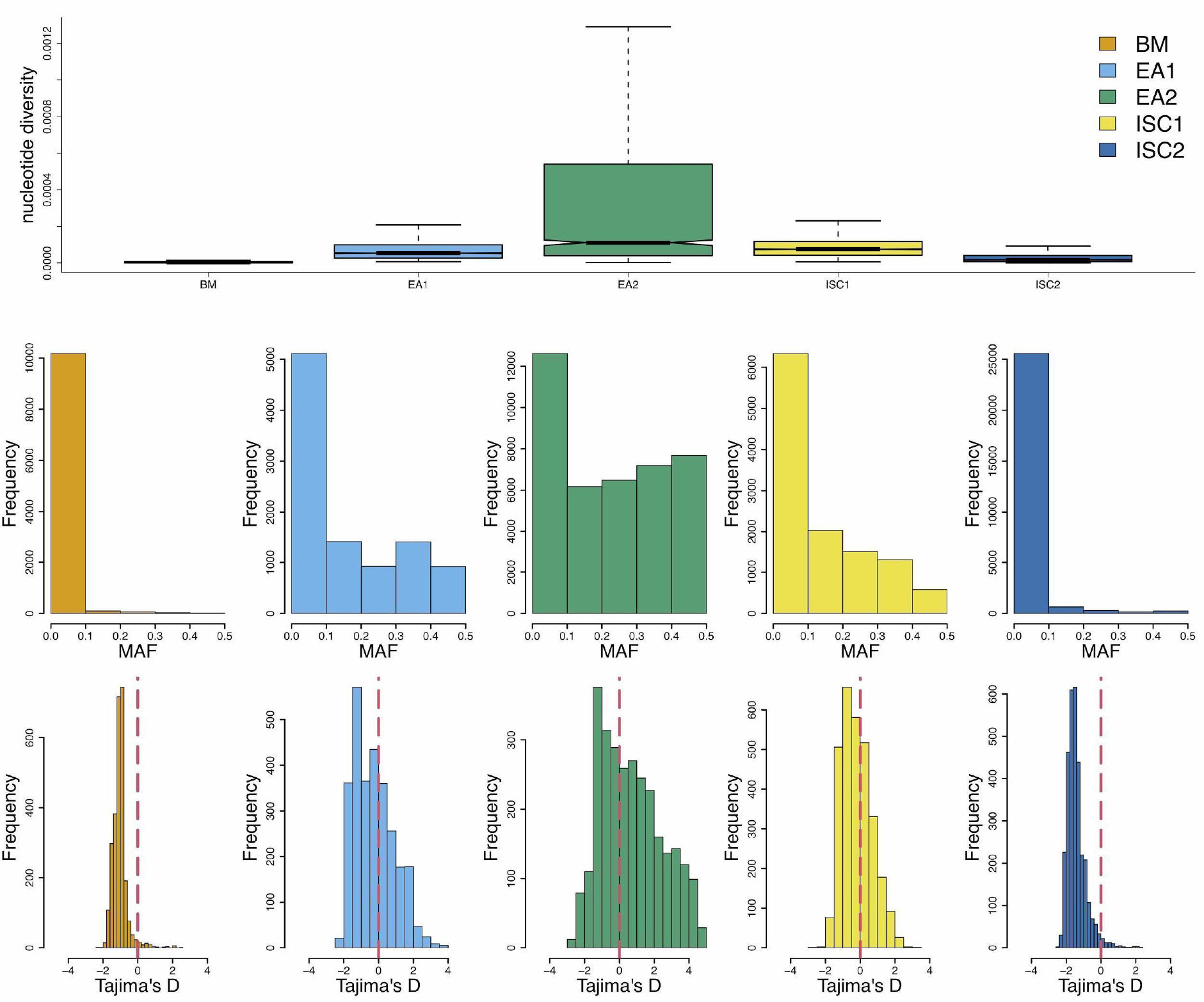
Population genetic statistics. Upper panel: nucleotide diversity (π); middle panel: Minor allele frequency (MAF); Lower panel: Tajima’s *D*.

As well as the very low diversity, the Brazil-Mediterranean and smaller East African (EA2) population contain more indel polymorphisms (Table 1), with a SNP:indel ratio of 6:1 and 5:1, respectively, compared to EA1 (11:1), ISC1 (10:1) and ISC2 (9:1), as expected for eukaryote polymorphisms, where SNPs typically outnumber indels (Mullaney et al. 2010; Jeffares et al. 2015). Extensive variant-calling quality control showed that this excess of indels is unlikely to be artefactual (Supplementary Figure 4). It is possible that this is due to the accumulation of weakly-deleterious indel alleles when the Brazil population was established from European *L. infantum* populations (Boité et al. 2019).

### Analysis of *L. donovani* species complex populations

Genetic diversity summary statistics vary considerably between populations (Figure 2, Table 1), consistent with these varied demographic histories in different locations. For example, the initial population within the ISC (ISC1) has a normally-distributed Tajimas’s *D*, whereas the Tajimas’s *D* is strongly skewed to negative values in the emerging ISC2 population. To select groups of strains that will approximate panmictic populations, we used the ADMIXTURE analysis to identify isolates that were confidently assigned to one population (*n*=433), rather than being inter-population hybrids. This selection resulted in 59 isolates from East Africa (two populations of 41 and 18), 226 from the ISC (two populations of 211 and 15) and 127 from Brazil-Mediterranean (Table 1). To characterise BS in these populations we applied the *NCD2* test (Bitarello et al. 2018) and *Betascan2* test (Siewert and Voight 2020) in 10 kb windows to each of the five populations.

### Are targets of balancing selection shared between populations?

In some circumstances balancing selection can be maintained as populations or species diverge (Siewert and Voight 2017; Bitarello et al. 2018; Mérot et al. 2020; Wang et al. 2020; Ding et al. 2021). Given these examples, we examined whether any genes had maintained BS between populations of the *L. donovani* species complex, indicating long term BS. To assess this without relying on shared polymorphisms, we used the *Betascan2* maximum and *NCD2* minimum scores for each gene, for each population as a summary statistic (see methods) as tests were consistent (Supplementary Figure 12). We find scant evidence for shared BS from *Betascan2* scores. We define *Betascan2* outlier genes as those in the upper 5% of *Betascan2* scores for their population. There was little overlap in these outliers; 701 genes are outliers in at least one population, only 42 of these (6%) were outliers in two or more populations (**Supplementary Table 3**) and only 9 are outliers in three or more populations (1%). The *NCD2* metric identified more overlap between populations, but long term BS is still the exception; 1627 genes were 5% outliers in at least one population and only 195 (11%) were outliers in more than one. However, since the *NCD2* metric measures the similarity of allele frequencies to a target frequency (0.5 in our case; Bitarello et al. 2018), genes that are merely subject to weaker purifying selection will have elevated *NCD2* scores.

Another possibility is that weak polygenic BS that operates on a number of genes, perhaps transiently. This may be the case for frequency-dependent BS, for example in exported and cell surface-located erythrocyte membrane and exported proteins in *Plasmodium falciparum* (Volkman et al. 2002; Jeffares et al. 2007; Claessens et al. 2014). In this scenario, we might expect BS targets in one population to predict genes with higher metrics in other populations, due to a history of weak BS. We examine this using only *Betascan2,* because we suspect that the measure of correlated allele frequency that Betascan ultises will be less confounded by weak purifying selection. Again, we find little evidence that weak BS acts consistently between populations. For example, genes that are 5th percentile *Betascan2* outliers from the East African EA1 population, do not have significantly elevated *Betascan2* scores in any other population. After comparing *Betascan2* outliers in EA1, ISC1 and the Brazil-Mediteranean population (BM) to all other populations, the only strongly significant enrichment between was between the two populations from the Indian subcontinent (ISC1 outliers are predictive of higher scores in ISC2, *P* = 2 x 10^-4^, Supplementary Figure 18). Since the ISC2 population is derived from ISC1 relatively recently, we can expect some aspects of the genetic diversity to be maintained. Hence, signals of balancing selection in this species complex are generally restricted to one population.

### Identifying genes that are subject to balancing selection

To advance research in *Leishmania* it would be useful to identify the most likely targets of BS. As a pragmatic approach, we sought to identify genes with robust and strong signatures from multiple metrics. To achieve this, we selected genomic windows that were in the 1st or 99th percentile of either the *NCD2* test or *Betascan* tests, respectively. To identify the genes within *NCD2*/*Betascan* windows that are likely targets, we calculated nucleotide diversity (π) and Tajima’s *D* (Tajima 1989) for each gene, and selected genes in the 90th percentile of either statistic as well-supported plausible targets. We then selected genes that were outliers in both categories (*NCD2 or Betascan2* and π or Tajima’s *D*). This intersection identified 33 genes (**Supplementary Table 4**). We manually vetted these to remove ‘hitchhikers’, genes whose high diversity was likely due to their proximity to a BS ‘driver’ gene. We also removed genes with suspicious read coverage, since gene duplications produce strong artefactual signals of BS (Supplementary Figure 13). Due to the stringent process of filtering, this method is not guaranteed to have equal power to detect BS in the five populations we examine, which are represented by different numbers of isolates and have patterns of genetic diversity (Figure 2, Table 1).

This screen identified 19 vetted candidate genes (Table 2, justification for vetting is genes in **Supplementary Table 4**). Candidate genes in the EA1 population, where the most were discovered, have nucleotide diversity that is 39-fold higher than the genome-wide median (Figure 3). Diversity is elevated in genome regions surrounding these target genes, and remains significantly elevated up to 130 kb from the targets. Since the mean size for chromosomes in *L. donovani* is 900 kb, this increase in diversity influences a large proportion of the genome. Furthermore, BS candidates are enriched for high MAF co-segregating sites (Supplementary Figure 14).

**Table 2:**
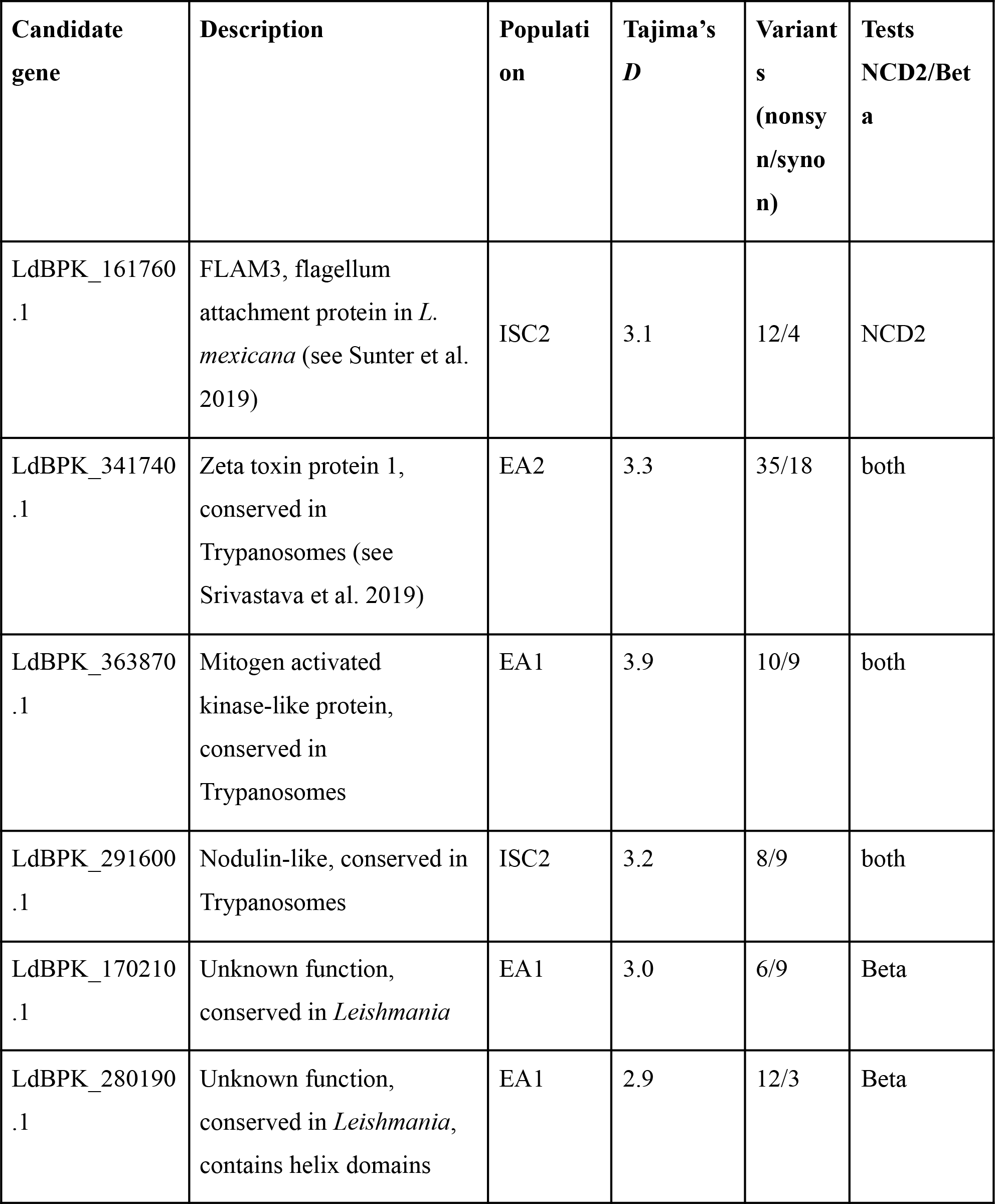
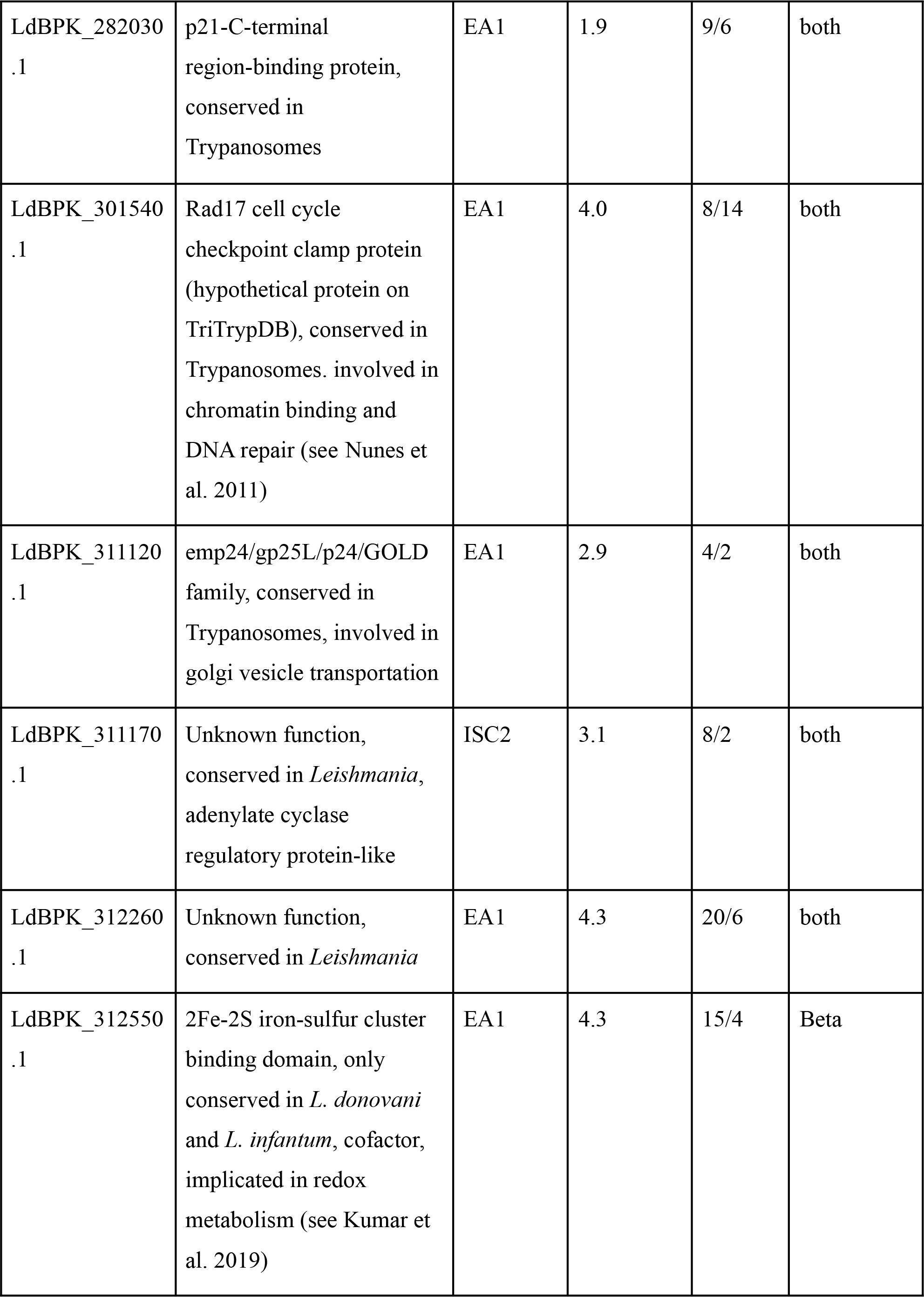
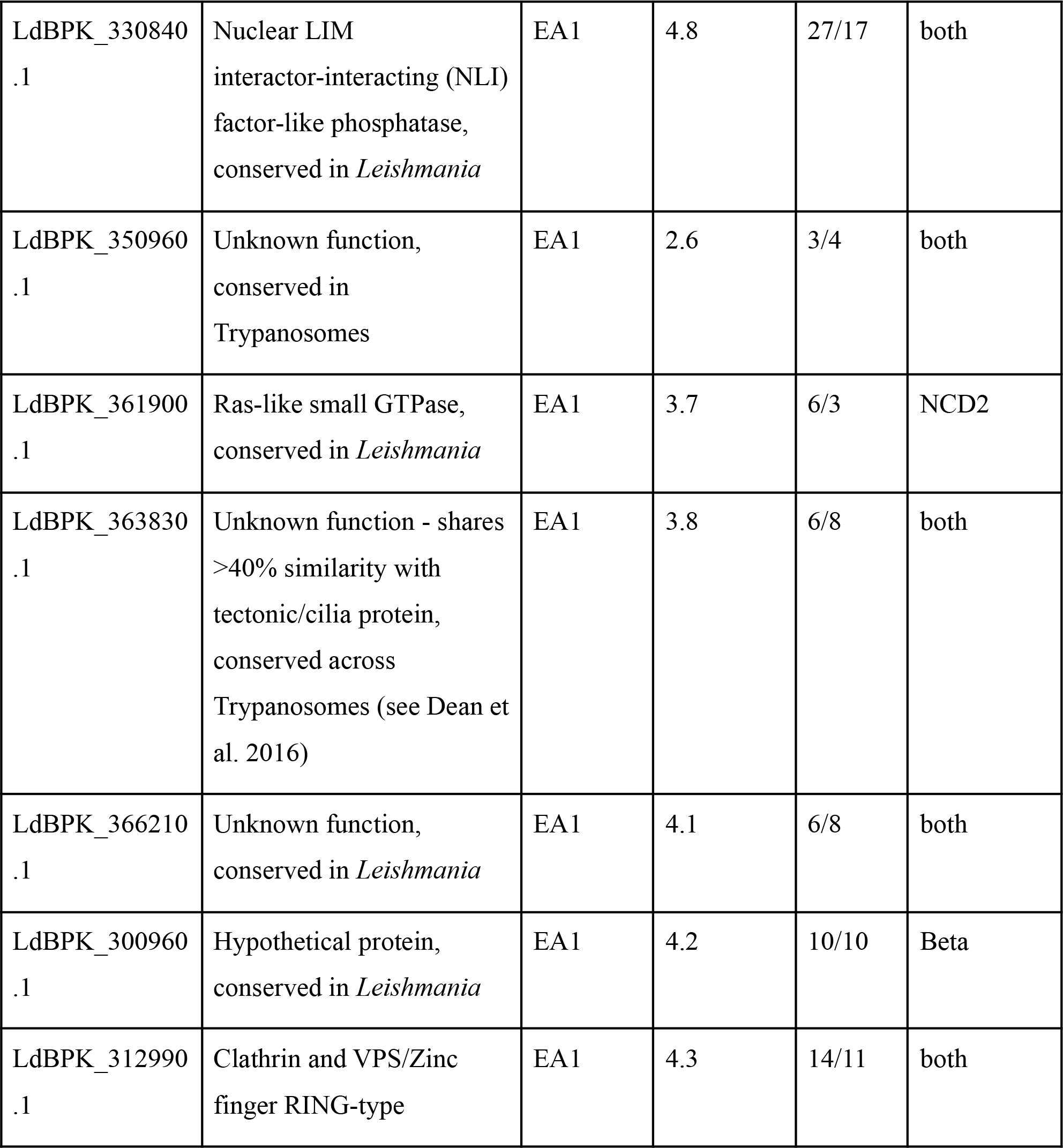
Candidates for gene subject to balancing selection in the L. donovani complex. Details of the method are described in Supplementary Figure 6 and **Supplementary Text 2**. All comparisons of tests for each population are available as Supplementary Figures 7-11, respectively. Where protein function is ‘unknown’ on TriTrypDB, we subjected each protein to BLASTp searches to obtain homology to other known proteins and ascertain conservation across trypanosomes.

**Figure 3.**
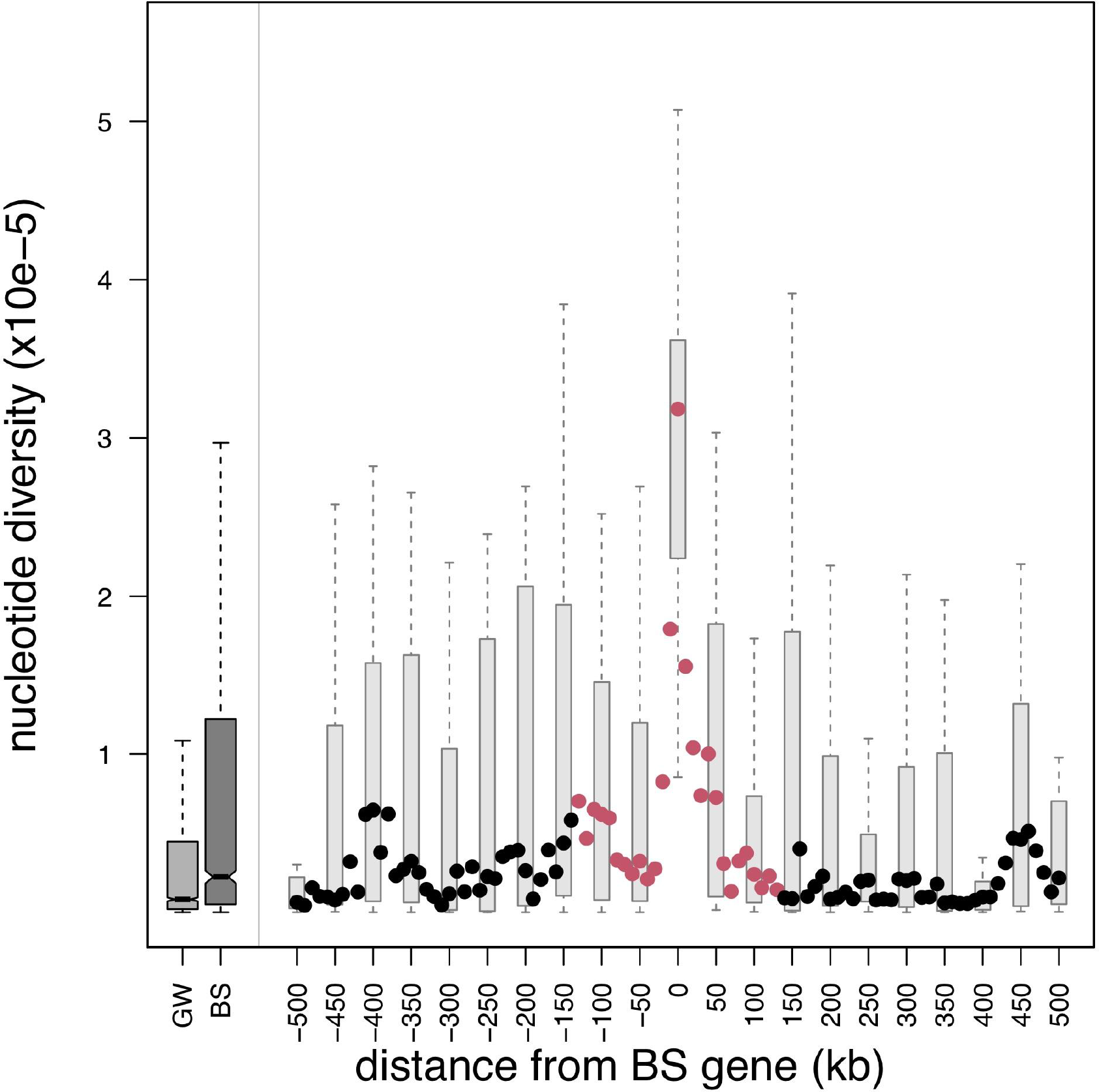
Diversity is significantly elevated in BS target regions. On the left we show the distribution of nucleotide diversity (π) genome-wide for the EA1 population (GW) and the distribution for the 500kb around all the 24 BS targets discovered in the EA1 population (including those that did not pass manual vetting)(BS). On the right, the filled circles show the median π (for all BS targets) every 10 kb up and downstream to 500 kb from the targets. Circles are red where the diversity at this distance is significantly higher than the genome-wide distribution and black otherwise (Wilcoxon signed rank tests < 1.5 x 10^-4^, using both up- and down-stream π values). The distribution of nucleotide diversity values for target genes is shown using box and whisker plots at 50kb intervals. Consistent with the lack of evidence for shared BS between populations, the genes that are BS candidates in the EA1 population do not show significantly elevated Tajma’s *D* values in any other population (Supplementary Figure 17).

### Candidate genes for balancing selection in the *L. donovani* complex

Our manual vetting of BS candidates retained 19 genes (Table 2), of which 15 were discovered in the EA1 population. We did not discover any reliable candidates in the two populations that appear to be expanding following a bottleneck (BM and ISC1), nor in the population consisting entirely of *L. infantum* isolates (BM). Figure 4 illustrates the variety of robust genetic signatures that implicate four of these genes. All vetted genes contain similarly robust signatures (**Supplementary Figure 16**).

**Figure 4.**
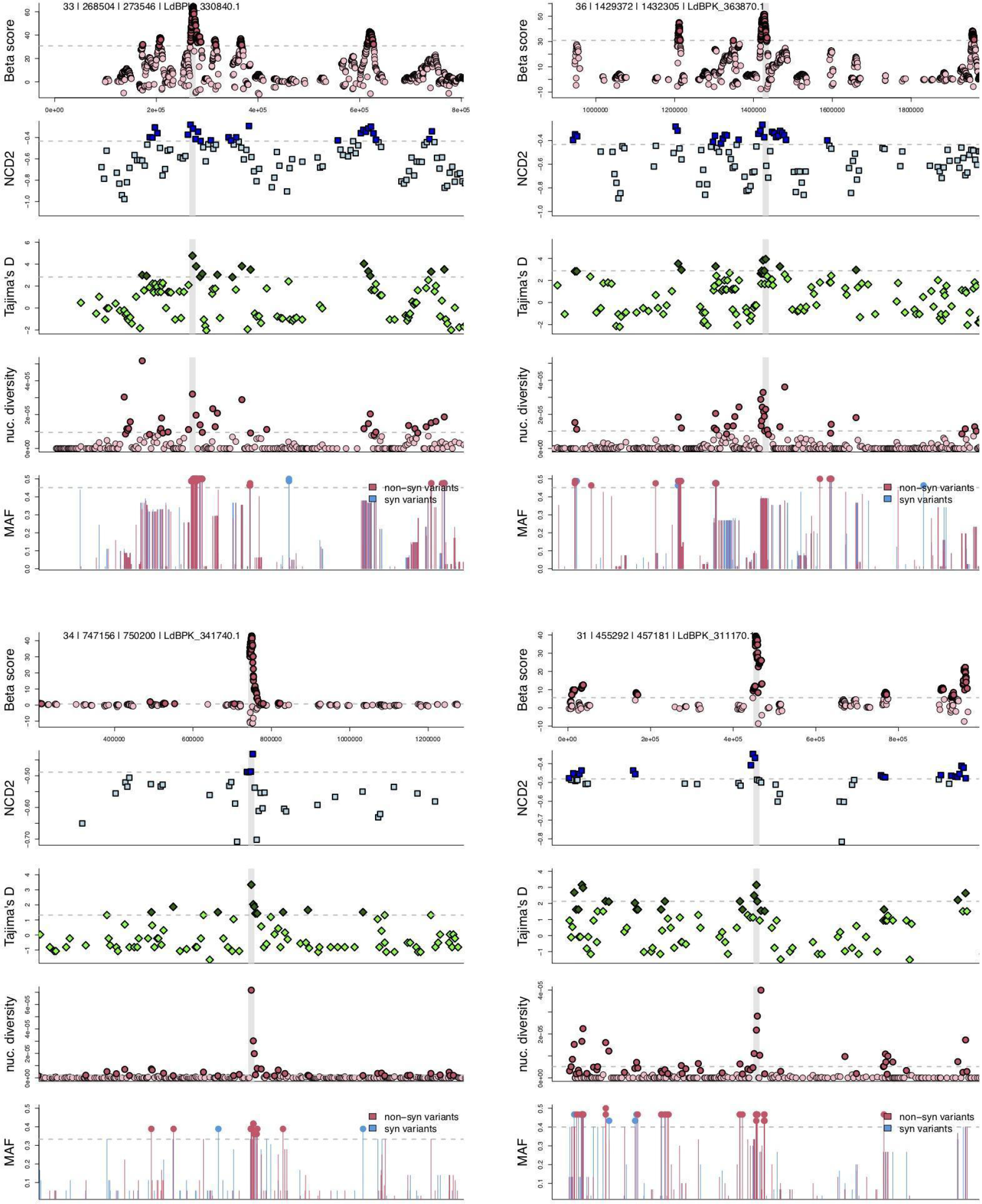
Candidate genes show multiple genetic signatures of balancing selection. We show *Betascan2*, *NCD2*, Tajima’s *D*, nucleotide diversity (π), and minor allele frequency (MAF) in a 500 kb window around four candidate genes. The location of the candidate gene is indicated by a vertical grey bar. The population-specific 90th percentile for each metric is shown as a horizontal dashed line, scores that are above this are drawn in darker shades, or plotted with a filled dot for MAF. Panel titles indicate the chromosome, gene start and end coordinates, and gene ID. Genes and populations where BS detected are; top left NLI interacting factor-like phosphatase-like phosphatase LdBPK_330840.1 (EA1), top right mitogen activated kinase-like protein LdBPK_363870.1 (EA1), bottom left putative Zeta toxin LdBPK_341740.1 (EA2), bottom right hypothetical protein LdBPK_311170.1 (ISC2). Similar plots are shown for all candidate genes in **Supplementary Figure 16**.

Several genes caught our attention as interesting targets of selection. LdBPK_291600.1 encodes a transmembrane protein containing a nodulin-like domain. Such proteins have been implicated in membrane transport and iron homeostasis (Laranjeira-Silva et al. 2018) in *Leishmania*.

The zeta toxin domain protein (LdBPK_341740.1, Figure 4) is indicated as a BS target by both *Betascan2* and *NCD2* metrics in the East African population EA2. The gene also has high nucleotide diversity in EA1 (**Supplementary Table 5**). The zeta domain is positioned at 744-861aa, with two non-synonymous variants resulting in the changes Leu747Thr and Ser752Phe, respectively, from the reference genome. The zeta toxin is part of the Type-II toxin-antitoxin (TA) module identified in prokaryotes, with homologues only recently discovered in *Leishmania* (Srivastava et al. 2019). The toxin component of the TA module acts against cellular processes such as translation, and is neutralised by the antitoxin component in favourable conditions. Sharing similar functional domains and activity with the *E. coli* homologue (Srivastava et al. 2019), *L. donovani* zeta toxin may therefore also be implicated in stress response and/or virulence (Rocker and Meinhart 2015).

The *FLAM3* gene (LdBPK_161760.1) encodes a flagellum attachment zone (FAZ) protein essential for host interaction (Sunter et al. 2015; An and Li 2018; Sunter et al. 2019). Our strict criteria has indicated this gene as under BS only in the Indian subcontinent population ISC2, but it also has high nucleotide diversity in four of the five populations (**Supplementary Table 5**). The *FLAM3* protein contains a clustered mitochondria (CLU) domain and a domain of repeats (Sunter et al. 2015). The majority of variants in ISC2 occur between these domains, with none falling within the CLU domain (**Supplementary Table 6**).

LdBPK_363870.1 (Figure 4) encodes a protein with similarity to mitogen-activated protein (MAP) kinases, a large family of enzymes implicated in antimony drug resistance (Ashutosh et al. 2012; Raj et al. 2019). While this gene was classified as a BS outlier only in the East African population EA1, its nucleotide diversity was also high in ISC1,2 and EA2 populations (**Supplementary Table 5**). LdBPK_363870.1 is also an orthologue of *L. mexicana* LmxM.36.3680, and therefore an endomembrane protein of the STE family (Baker et al. 2020).

Two candidate genes, LdBPK_311120.1 and LdBPK_361900.1, encoding a member of the emp24/gp25L/p24/GOLD family and a ras-like GTPase, respectively, are implicated in protein trafficking and export (Anantharaman and Aravind 2002; Field 2005). Other than this commonality, we do not observe any strong signals of functional enrichment in the 19 BS targets. There were no significant enrichments for gene ontology containing more than one gene and no significant enrichment for proteins with transmembrane domains or predicted signal peptides, or any enrichment in particular regions of the genome.

## Discussion

Here we sequenced 93 strains of *L. infantum* from Brazil, contributing to a worldwide collection of *L. donovani* complex isolates along with previous analyses (Imamura et al. 2016; Carnielli et al. 2018; Zackay et al. 2018; Franssen et al. 2020). Our analysis of this population is consistent with these previous studies, showing that the *L. infantum* population in Brazil contains very little genetic diversity (Carvalho et al. 2020; Schwabl et al. 2021). A consistent observation in analysis of this species complex is that populations from East Africa, India and Brazil are substantially genetically differentiated, a result that we reiterate here (Figure 1). In this study of balancing selection, we also show that signals of selection largely differ between populations.

Relatively few studies have attempted balancing selection screenings in parasites (reviewed by (Weedall and Conway 2010). Our study uses the *Betacan2* (Siewert and Voight 2020) and *NCD2 (Bitarello et al. 2018)* metrics that have been developed recently and have been shown outperform classic metrics under models of BS, such as Tajima’s *D* (Tajima 1989). These analytic tools, and the use of multiple populations should produce an analysis at least as sensitive as previous screens for balancing selection searches within parasites such as *Plasmodium* (Tetteh et al. 2009, 6 genes; Amambua-Ngwa et al. 2012, 25 genes; Nygaard et al. 2010, 19 genes).

These candidate genes possess varying functions within the parasite. The flagellum attachment gene *FLAM3* is a striking candidate in view of the importance of the encoded protein in parasite cellular structure, proliferation and differentiation (Sunter et al. 2019). Furthermore, candidate genes LdBPK_311120.1 and LdBPK_361900.1, which encode a member of the emp24/gp25L/p24/GOLD family and a ras-like GTPase, respectively, may influence trafficking of virulence factors, and subsequently interaction with their host. Given the lack of detailed studies of these candidates in the *L. donovani* species complex, studies of the cell biology of these proteins will be useful next steps.

Our analysis suggests that the performance of *Betascan2* and *NCD2* BS tests are dependent on population demography and history, despite claims that the tests are robust to demography (Siewert and Voight 2017; Bitarello et al. 2018; Siewert and Voight 2020; Stern and Lee 2020). This is likely due to our pragmatic screening criteria, that required >5 variants per genomic window (*Betascan2, NCD2*) and >5 variants per gene (Tajima’s *D* and π metrics). The low-diversity Brazil and ISC1 populations contain far fewer regions that satisfy these criteria. We expect this is because the population bottlenecks experienced by these populations has reduced its genetic diversity, ablating strong BS signatures. To our knowledge, this has not been reported in other species. Only two populations (EA1 and ISC2) appear to have relatively stable population sizes and sufficient nucleotide diversity to identify BS candidates using our pragmatic methods (Table 1). More complex methods that employ population models to detect BS are available (DeGiorgio et al. 2014; Cheng and DeGiorgio 2019; Cheng and DeGiorgio 2020). We chose not to employ these because we do not believe that *L. donovani* complex populations are sufficiently understood to be meaningfully modelled at this stage. For example, there is no accurate estimate of the recombination rates or mutation rate in these species. Our approach was to divide the samples into multiple populations and exclude potential between-population hybrids, with the expectation that this would alleviate some of the issues resulting from population structure.

It is possible that other evolutionary changes caused some of the signals we observe, including introgression events, partial selective sweeps or transient heterozygosity excess that can occur as a consequence of adaptation (Sellis et al. 2011). Local adaptation can also lead to an appearance of within-population diversity and/or excess heterozygosity (Wood et al. 2008; Eizaguirre and Lenz 2010; Ellison et al. 2011; Keller et al. 2011), particularly when the distribution of local ‘niches’ is not well understood, which is generally the case with *Leishmania*. It is possible, for example, that multiple variants exist within these genes as adaptations to regional differences in host or sandfly cellular/extracellular environments. Parasite genes may vary with regional variations in HLA loci that affect susceptibility to visceral leishmaniasis (Blackwell et al. 2020). In any case, the candidate genes we identify warrant further study.

It is possible that balancing selection of single-copy genes is not the most important mechanism that maintains diversity within the *L. donovani* complex, or protozoan parasites generally. The effects of multi-copy gene families encoding RIFIN, STEVOR and PfEMP1 variant surface antigens in pathogenesis of *Plasmodium* parasites are well-described, for example (Wahlgren et al. 2017). These genes are typically removed from BS screens, because multi-copy genes will produce artefactual signals of BS when analysed with current bioinformatics methods (Supplementary Figure 15). While variants in duplicated regions or multiple copies of genes may allow the parasite to maintain diversity, it is an open question whether this diversity is maintained by neutral processes or balancing selection.

In summary, our description of diversity in the *L. donovani* species complex provides insight into the global populations of this parasite. We show that these populations are genetically divergent, with largely independent signals of balancing selection. Our discovery of a handful of genes with robust signatures of balancing selection provides candidate genes for the study of host-parasite and host-vector interactions.

## Materials and Methods

### Ethics

Samples from Brazil were obtained as part of a broad study for genomic studies in the Laboratory of Leishmaniasis at the Institute of Tropical Medicine Natan Portella, approved by the Research Ethics Committee of the Federal University of Piauí (approval ID number 0116/2005). All methods were performed according to the approved guidelines and regulations. A written informed consent was obtained from all study participants or their legal guardians.

### Strain culture and genome sequencing

Bone marrow aspirates were obtained from the routine diagnosis of patients admitted to the Natan Portella Tropical Diseases Institute in Teresina-PI, Brazil. Aspirates were inoculated into a mixed culture medium NNN (Neal, Novy, Nicolle) containing 2 ml of Schneider’s medium supplemented with 10% fetal bovine serum, 2% urine and penicillin 10,000 U/ml, and streptomycin 10 mg/ml. The positive isolates in mixed media were expanded in Schneider’s liquid medium under the same conditions mentioned above. Extraction of DNA from the parasites was performed after washing to remove culture medium, using Qiagen Blood and Tissue kit was used according to the manufacturer’s recommendation.

Genome sequencing was performed on Illumina HiSeq 2500 machines (or similar) to produce paired end 150 nt reads. The majority (95%) of the samples were sequenced to provide mapped read coverage of ≥ 30x (mean 97x, minimum 19x). Raw sequencing reads were submitted to NCBI’s sequencing read archive (SRA) under the BioProject accession PRJNA702997 (note to reviewers: data are scheduled to be released upon publication).

### Sequence analysis/variant calling

Publicly available *L. donovani* complex data was downloaded in FASTQ format from the European Nucleotide Archive (ENA: https://www.ebi.ac.uk/ena). Full list of strain names/ENA numbers in **Supplementary Table 1**. The *L. donovani* reference genome (strain BPK282A1) was downloaded from TriTrypDB (version Nov 2019). Strain reads were mapped to the reference using bwa v.0.7.17 (Li and Durbin 2009), converted to bam, sorted, indexed and duplicates removed with SAMtools v.1.9 (Li et al. 2009). For each strain, SNPs and indels were called using The Genome Analysis Toolkit (GATK) HaplotypeCaller v.4.1.0.0 (Depristo et al. 2011) using the ‘discovery’ genotyping mode Freebayes v.1.3.2 (https://github.com/ekg/freebayes) accepting calls with a minimum alternative allele read count ≥ 5. We accepted calls discovered by both methods, merged all VCFs and regenotyped with Freebayes. The regenotyped VCF was sorted with Picard SortVcf (https://broadinstitute.github.io/picard/) and indexed with GATK IndexFeatureFile. SNP hard-filtering was performed with BCFtools (https://samtools.github.io/bcftools/) on biallelic variants only, to remove sites with any of the following: DPRA < 0.73 or > 1.48; QA or QR < 100; SRP or SAP > 2000; RPP or RPPR > 3484; PAIRED or PAIRPAIREDR < 0.8; MQM or MQMR < 40. As chromosome 31 is generally supernumerary, we specified DPB < 30401 or > 121603 to be removed, and for remaining chromosomes, DPB < 18299 or > 73197 (<0.5x or >2x median DPB). Biallelic indels were filtered to remove sites with any of the following: DPRA < 0.73 or > 1.48; QA or QR < 100; SRP or SAP > 2000. VCF annotation was performed with the snpEff v.4.3 package (Cingolani et al. 2012) using the default Leishmania_donovani_BPK282A1 database included with the software. SnpSift filter with the option “ANN[*].EFFECT has ‘missense_variant’” was used to extract nonsynonymous sites.

With this variant filtering we observed a correlation between minor allele frequency (MAF) and read coverage at SNP sites (Supplementary Figure 13). Modelling showed that duplications resulted in a systematic bias against calling rare alleles. We therefore removed any SNP/indel sites where the mean variant coverage within the ADMIXTURE-defined population was ≥1.5x larger than the median coverage (corresponding to triploid sites in a generally diploid chromosome), or ≥1.25x larger than the median coverage for chromosome 31 (corresponding to tetraploid sites in a generally triploid chromosome). We also removed sites where coverage was highly variable, by excluding sites in the upper 5th percentile of the coverage standard deviation. In each population this filtered ∼5-7% of sites. Mapping coverage was ascertained by SAMtools bedcov for each gene in the multipopulation VCF. After this filtering, the correlation between MAF and read coverage was either far less significant or removed completely. This filtering retained 10377 out of a possible 10778 sites in population ISC1; 9781 out of a possible 10227 sites in population ISC2; 40127 out of a possible 41957 sites in EA1; 11757 out of a possible 12365 sites in EA2 and 26884 out of a possible 28281 sites in Brazil-Med.

To validate the variant filtering we produced a *de novo* assembly of the MHOM/BR/06/MA01A *L. infantum* isolate from Brazil (Carnielli et al. 2018), mapped Illumina reads from the same isolate to the assembly, and called SNPs and indels as above. All calls should be heterozygous sites, or errors. Initial variant calling identified 4 SNPs and 23 indels, after filtering no SNPs or indels remained, consistent with a very low false positive call per strain. The MHOM/BR/06/MA01A *de novo* assembly will be described elsewhere. Briefly, the assembly was produced using Oxford Nanopore Technology (ONT) reads to 110x coverage, assembled with Canu v.1.9 (Koren et al. 2017), polished once using ONT reads using Nanopolish v.0.9.2 (Loman et al. 2015) and thrice with Illumina reads using Pilon v.1.22 (Walker et al. 2014).

### Phylogenetic analysis

VCF containing all variants from all 477 isolates was converted to PHYLIP format using vcf2phylip (available at https://github.com/edgardomortiz/vcf2phylip/tree/v2.0). This produced an alignment of 283378 sites. IQ-TREE v.1.5.5 (Nguyen et al. 2015) was used to perform Maximum Likelihood (ML) phylogenetic analysis with the model GTR+ASC, which includes ascertainment bias correction, with 1000 bootstrap replicates and 1000 UFBOOT (Hoang et al. 2018) approximations to produce ML support values. The resulting tree was visualised with Figtree v.1.4.4 (available at http://tree.bio.ed.ac.uk/). Treefiles are available in supplementary material.

### Population and diversity analysis

For all population analyses we utilised only biallelic SNPs, pruning linked sites (*r*^2^ > 0.5) in 2 kb windows with a step size of 1 with PLINK v.1.9 (Purcell et al. 2007) using the option --indep-pairwise 2kb 1 0.5. This produced 194351 SNPs from the initial 353301 (158950 variants removed). ADMIXTURE v.1.3 (Alexander and Lange 2011) was run with *K* = 1-12. Principal component coordinates were produced with PLINK v.1.9.

Prior to balancing selection tests performed on the five populations (EA1, EA2, ISC1, ISC2, BM), mixed ancestry strains were removed from population VCFs. Population-specific VCFs were filtered with VCFtools v.0.1.15 (Danecek et al. 2011) to remove sites that were fixed within a population (option --mac 1). Repeat regions (see below) were also filtered out of VCFs at this stage. Tajima’s *D*, π and MAF were calculated on unpruned variants using VCFtools. Tests for balancing selection used biallelic SNPs and indels from each population. Copy-number variant and duplicated genome regions were removed from this analysis, as these regions will produce biases in allele frequencies towards common alleles, producing artifactual signals of BS (**Supplementary Text 1**, Supplementary Figure 5). Variant calling for multi-copy regions was beyond the scope of this study.

Repeat regions were determined as follows. Intergenic coordinates in *L. donovani* were extracted from the annotation .gff, downloaded from TriTrypDB (version Nov 2019) with BEDtools v.2.27.1 (Quinlan and Hall 2010) complement with default parameters. Intergenic regions were then extracted from the genome using BEDtools getfasta. Repeat regions in *L. donovani* were identified by nBLASTing v.2.7.1 (Altschul et al. 1990) intergenic regions against the rest of the genome, removing redundant hits and those <200nt in length. Resulting coordinates were converted to bed format for filtering.

### Balancing selection tests

The NCD2 test used software provided by (Bitarello et al. 2018), using windows of 1, 5 and 10 kb with step sizes of 0.5, 2.5 and 5kb, respectively. 10 kb windows sizes were used in this study. A list of fixed differences between *L. donovani* populations (total 285 isolates) and the ‘pure’ Brazilian-Mediterranean *L. infantum* population (127 isolates) was used as the outgroup for analysis. Fixed differences were determined using bcftools isec called on VCFs of all *L. donovani* populations and *L. infantum* containing fixed variants, resulting in 76284 fixed difference sites. Resulting target frequencies (*tf*) and Equation 2 of (Bitarello et al. 2018) were used to generate *Ztf-IS* scores, with the exception of using the standard deviation (SD) for each number of informative sites (IS) rather than simulated SD. *P* values for each window were calculated by assigning a rank based on *Ztf-IS* score and dividing by the total number of scanned windows.

The BetaScan2 test (Siewert and Voight 2020), using default parameters, using the file format generated from the variants using glactools (Renaud 2017; available at: https://github.com/grenaud/glactools), using the Brazil-Mediterranean *L. infantum* population as the outgroup. We performed the test on each individual population. *Betascan2* and *NCD2* scores are calculated in windows around each variant site. To obtain values for each gene, we used the maximum *Betascan2* score for all variants within the gene and the minimum *NCD2* score within each gene (since low *NCD2* scores are indicators of BS).

### Gene Ontology analysis

Gene Ontology (GO) descriptions and gene details for the *L. donovani* BPK282A1 reference genome were downloaded from TriTrypDB. GO enrichment analysis was performed using the PANTHER service on tritrypdb.org. Proteins that were classed as ‘hypothetical’ or of ‘unknown function’ were BLASTed against the non-redundant protein sequences (nr) database of NCBI to obtain possible identity by shared homology, and to determine conservation across trypanosomes.

## Acknowledgements

This work has been produced as part of the UK:Brazil Joint Centre Partnership in Leishmaniasis. This project was undertaken on the Viking Cluster, which is a high performance computer facility provided by the University of York. We are grateful for computational support from the University of York High Performance Computing service, Viking and the Research Computing team. We acknowledge Shoumit Dey and Martina Stoycheva for contributions to the VCF pipeline, and João Cunha for comments on the manuscript. We acknowledge the support of John Davies for the Brazil *Leishmania infantum* assembly and The Genomics and Bioinformatics Laboratory at The University of York for their assistance.

## Author contributions

CAG - performed data curation, sequence analysis and variant calling, phylogenetics, population and diversity analysis, balancing selection tests, gene ontology analysis, contributed to writing the manuscript

SJF - contributed to data curation, sequence analysis and variant calling pipelines, data analysis and writing the manuscript

VC - performed isolate culture and DNA extraction for Brazil samples

AA - contributed to population and diversity analysis

HK - contributed to population and diversity analysis

YPC - contributed to data analysis

SJ - performed library preparation and sequencing of isolates from Brazil

DLC - contributed to isolate collection in Brazil

JCM - obtained funding, contributed to data analysis and writing the manuscript

CHNC - obtained funding, collected and cultured isolates from Brazil, contributed to the manuscript

DCJ - obtained funding, initiated the project, supervised student work, assisted with data analysis, wrote the manuscript

## Supplementary data

**Supplementary Table 1** lists all isolates used in this study, including source, location, and ENA accession numbers, including genome data from Brazil (produced in this work). All sequence data generated in this study have been deposited under the NCBI BioProject accession PRJNA702997 and will be released upon publication. All Betascan2, NCD2, Tajima’s *D* and nucleotide diversity metrics and Supplementary Tables are available from the Figshare repository https://figshare.com/account/home#/projects/94292 (Note to reviewers: this place holder will be replaced when the article is accepted).

## Supplementary Text

**Supplementary Text 1. Duplications distort allele frequencies**

Initial analysis of the allele frequencies of these populations showed a tail of high MAF alleles, that would be consistent with strong balancing selection in some sites. However, closer inspection of these data showed that all populations had a strong positive correlation between read coverage and MAF (Supplementary Figure 5). Very high copy regions, which included multiple duplicated genes, were significantly enriched for common alleles. This bias is most likely due to systematic under-calling of rare alleles in duplicated genes, combined with the appearance of ‘balanced’ allele frequencies caused by variants fixed in one duplication (Supplementary Figure 15). These processes will distort allele frequencies, and result in artefacts in many of the methods used to detect balancing selection, such as Tajima’s *D* (Tajima 1989) and *NCD2* (Bitarello et al. 2018), and *BetaScan/Betascan2* which utilise correlated alleles (Siewert and Voight 2017; Siewert and Voight 2020). As the missing rare alleles are generally newer, this process will also reduce population distances (F_ST_), an expectation of long term balancing selection (Charlesworth 2006).

**Supplementary Text 2. Full description of BS tests and justifications**

To identify the most likely candidates of BS in these populations we applied a search strategy that used genomic window analyses, followed by gene-centric analysis (Supplementary Figure 5). To identify genomic windows that contain signals consistent with BS we applied two tests, *BetaScan2* (Siewert and Voight 2020) and *NCD2 (Bitarello et al. 2018)* over 10 kb windows around each segregating site, for each population. *BetaScan2* detects regions with alleles ‘balanced’ at correlated frequencies and a deficit of substitutions compared to the outgroup, while the *NCD2* test detects regions with an excess of alleles near a target frequency (0.5 in our case). Selection of outlier regions with *Betascan2* and/or *NCD2* scores in any population identified 258 genes (13 were identified in more than one population).

For gene-centric analysis, we identified specific genes that may be BS targets within these windows, therefore we calculated nucleotide diversity (π) and Tajima’s *D* (Tajima 1989; Bitarello et al. 2018; Siewert and Voight 2020) for all genes in all populations (**Supplementary Table 5**). We selected genes in the 90th percentile of either statistic for any population as the most likely targets of balancing selection. Neither of these gene sets were significantly enriched for any Gene Ontology categories after multiple-test correction. Selection of genes with both genomic window tests (*NCD2* or *Betascan*) and gene-centric metrics (Tajima’s *D* or π) identified 33 genes (**Supplementary Table 4**).

## Supplementary Figures

**Supplementary Figure 1.**
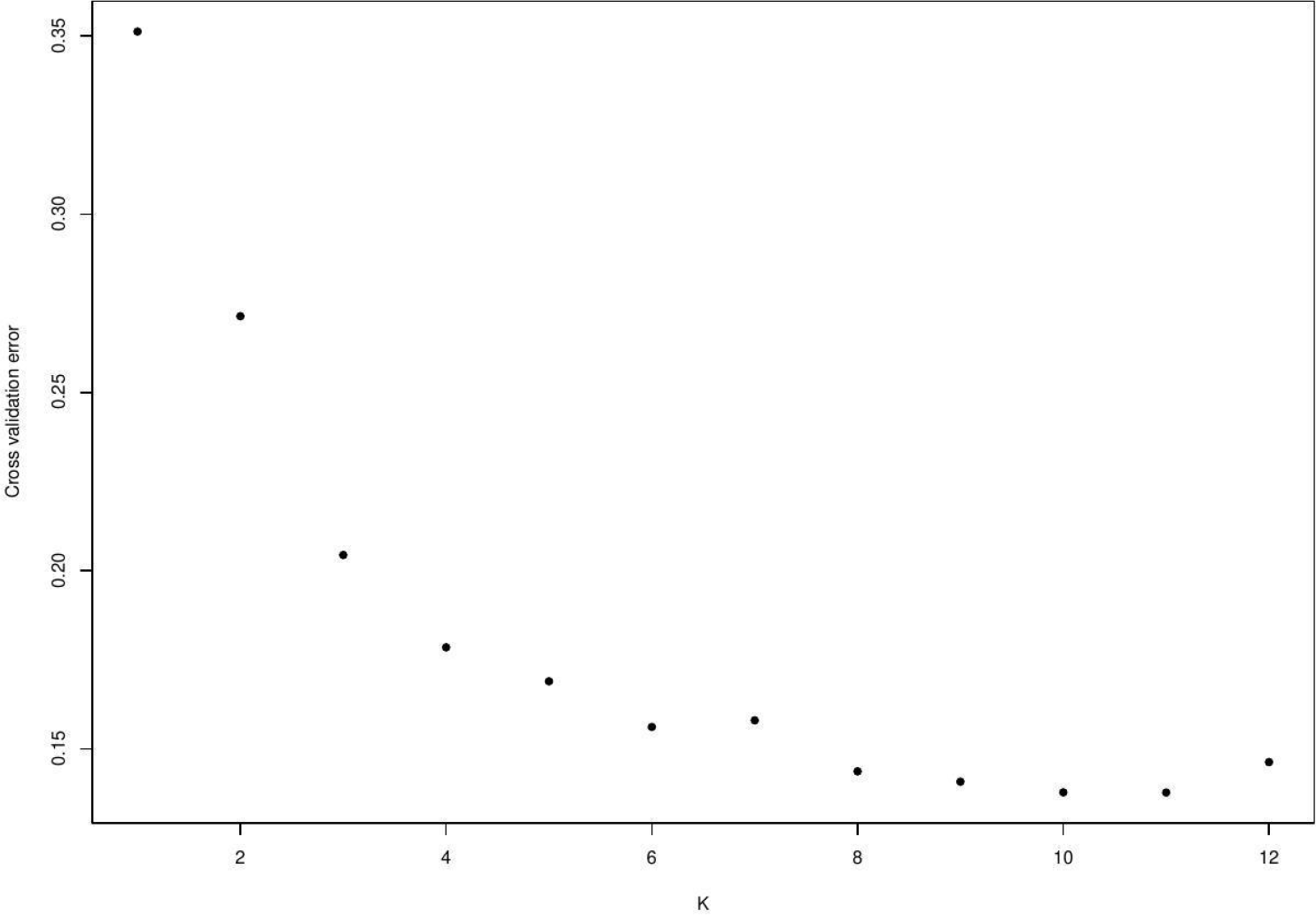
ADMIXTURE cross-validation error for *K*=1-12. ADMIXTURE analysis of 477 isolates indicated a true population number between 8-11.

**Supplementary Figure 2.**
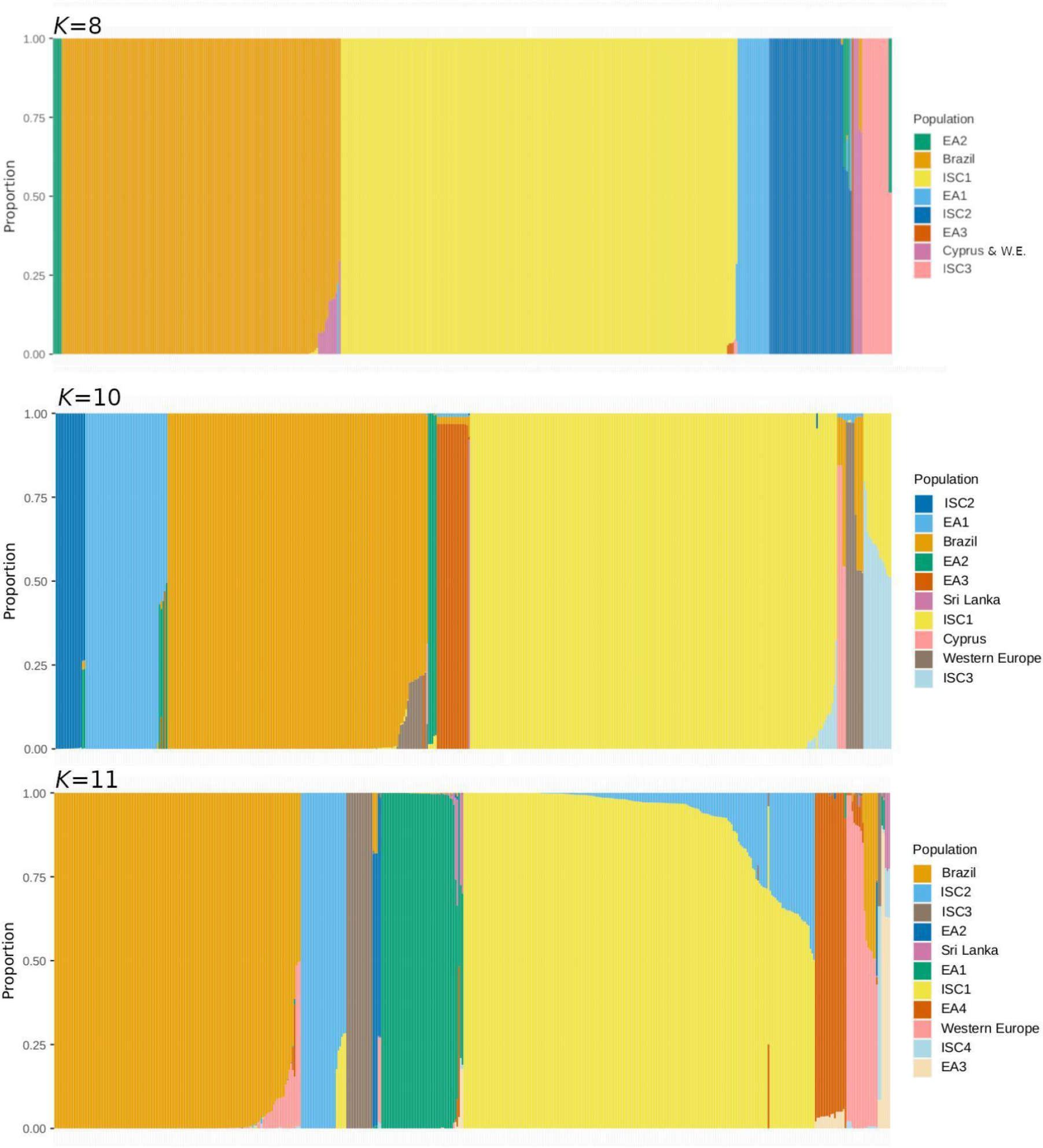
ADMIXTURE results for *K*=8, 10 and 11. Isolates assigned to populations when *K*= 8, 10 and 11. Each vertical bar represents a single isolate. We proceeded with *K*=9 (Figure 1) in line with previous analysis by (Franssen et al. 2020). Populations are named according to their predominant origin. WE = Western Europe.

**Supplementary Figure 3.**
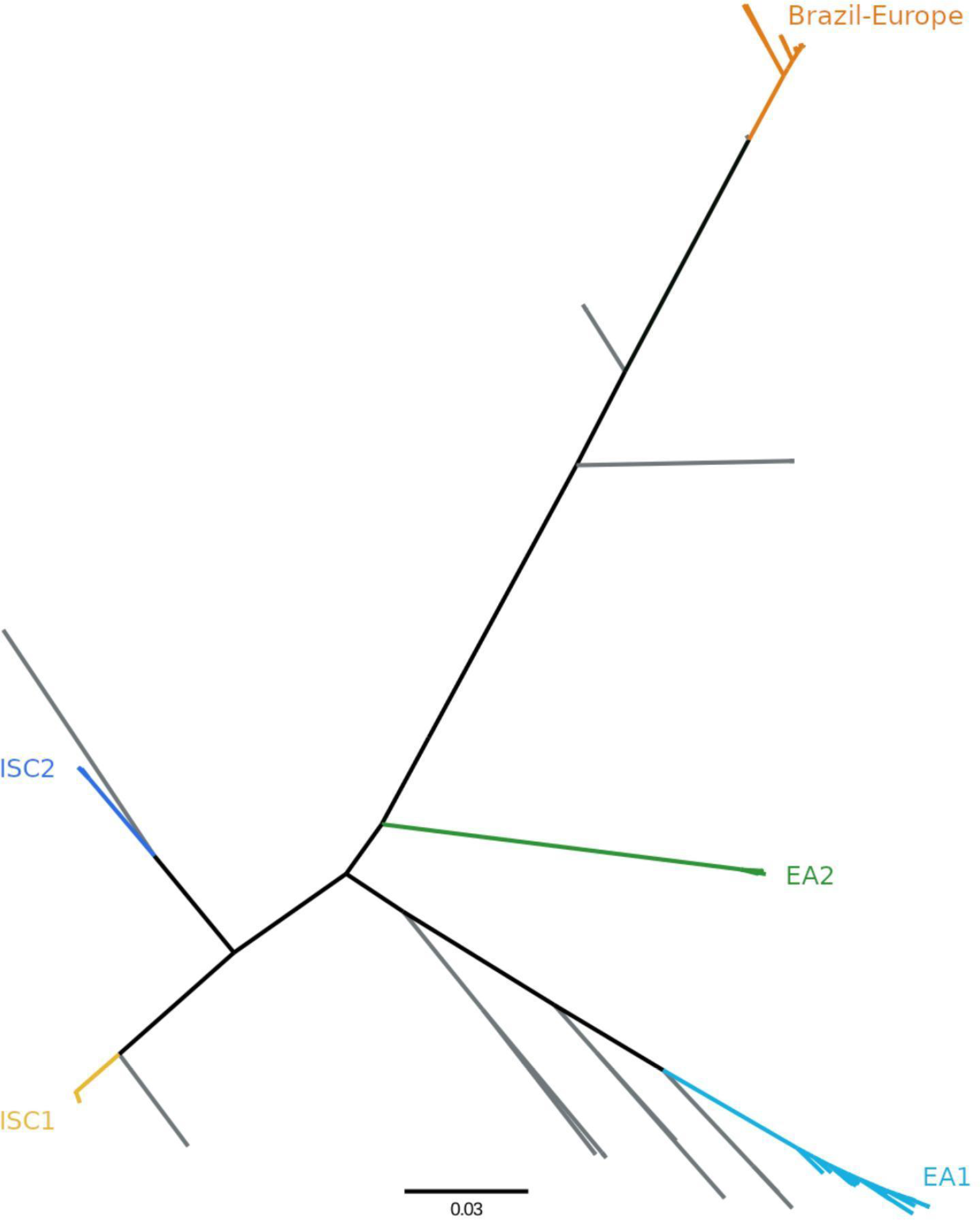
Unrooted phylogenetic tree of 477 isolates. Maximum likelihood phylogeny, based upon a SNP alignment of 477 sequences with 283378 variable sites. All visible branches are maximally supported (100% mlBP) unless indicated. The scale bar represents the number of nucleotide changes per site. Branches in grey are those isolates assigned to minor populations which we do not analyse further.

**Supplementary Figure 4.**
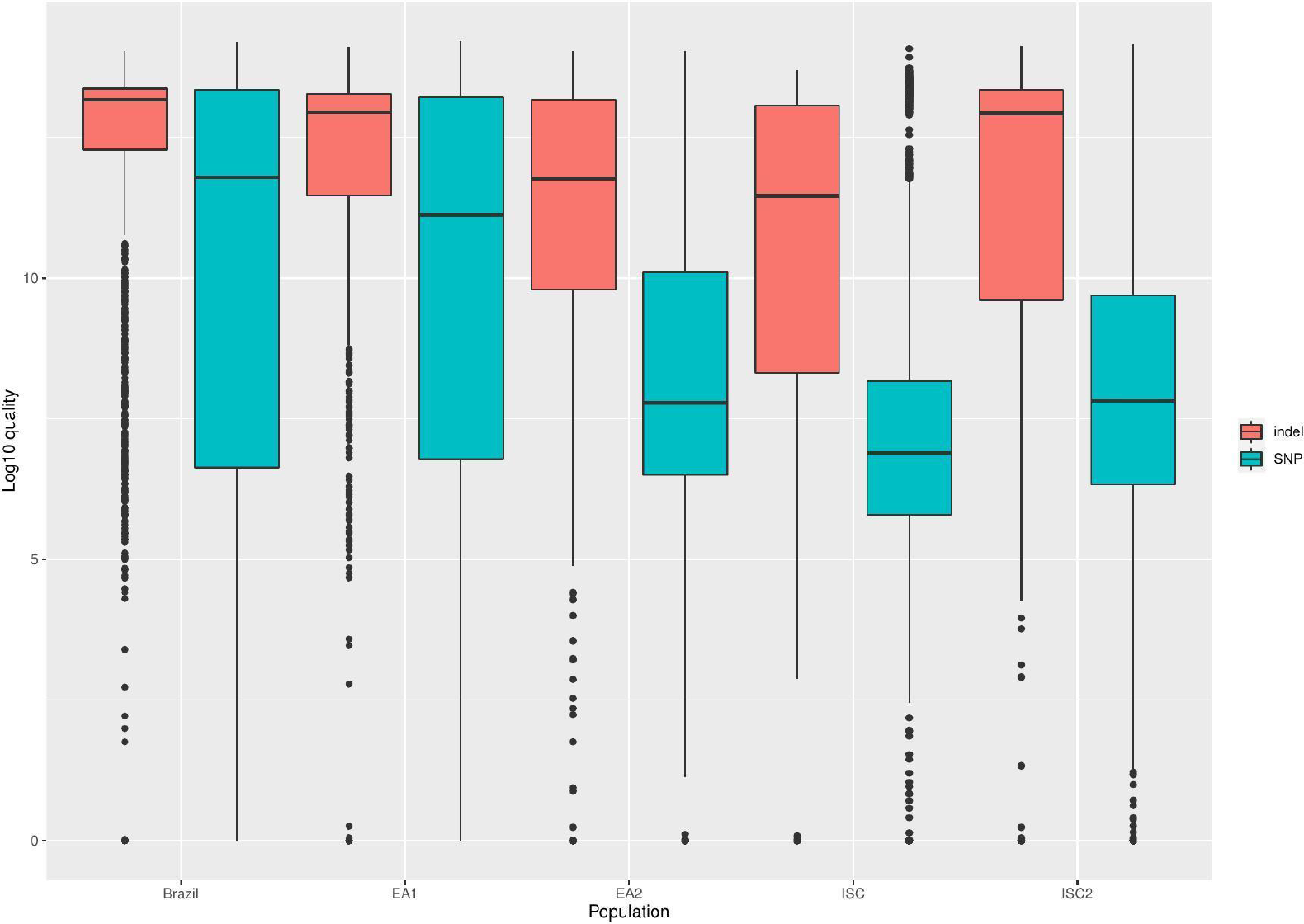
The excess of indels is unlikely to be artefactual. Log10 QUAL scores for SNP and indel positions are displayed for each population.

**Supplementary Figure 5.**
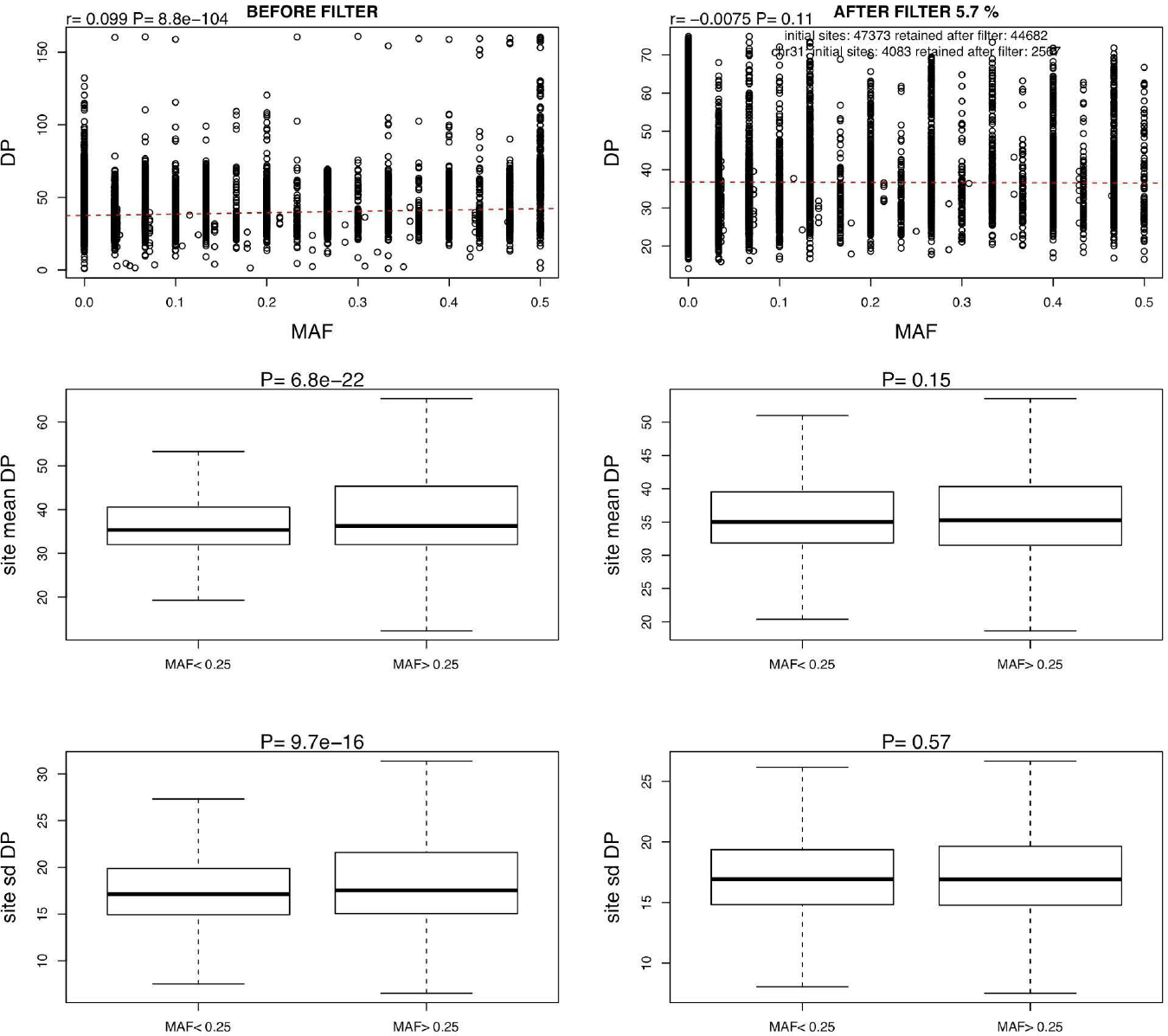
Duplications systematically bias allele frequencies. Data shown are from the ISC2 population. Left three panels show data prior to depth filtering. Top panel: a positive correlation between total read depth (DP) and minor allele frequency (MAF). Middle panel: sites with MAF > 0.25 are significantly higher read depth. Lower panel: sites with MAF > 0.25 are significantly higher variation in (standard deviation, sd) in read depth. Right three panels show the same presentations of data after read depth filtering. None of the relationships remain statistically significant.

**Supplementary Figure 6.**
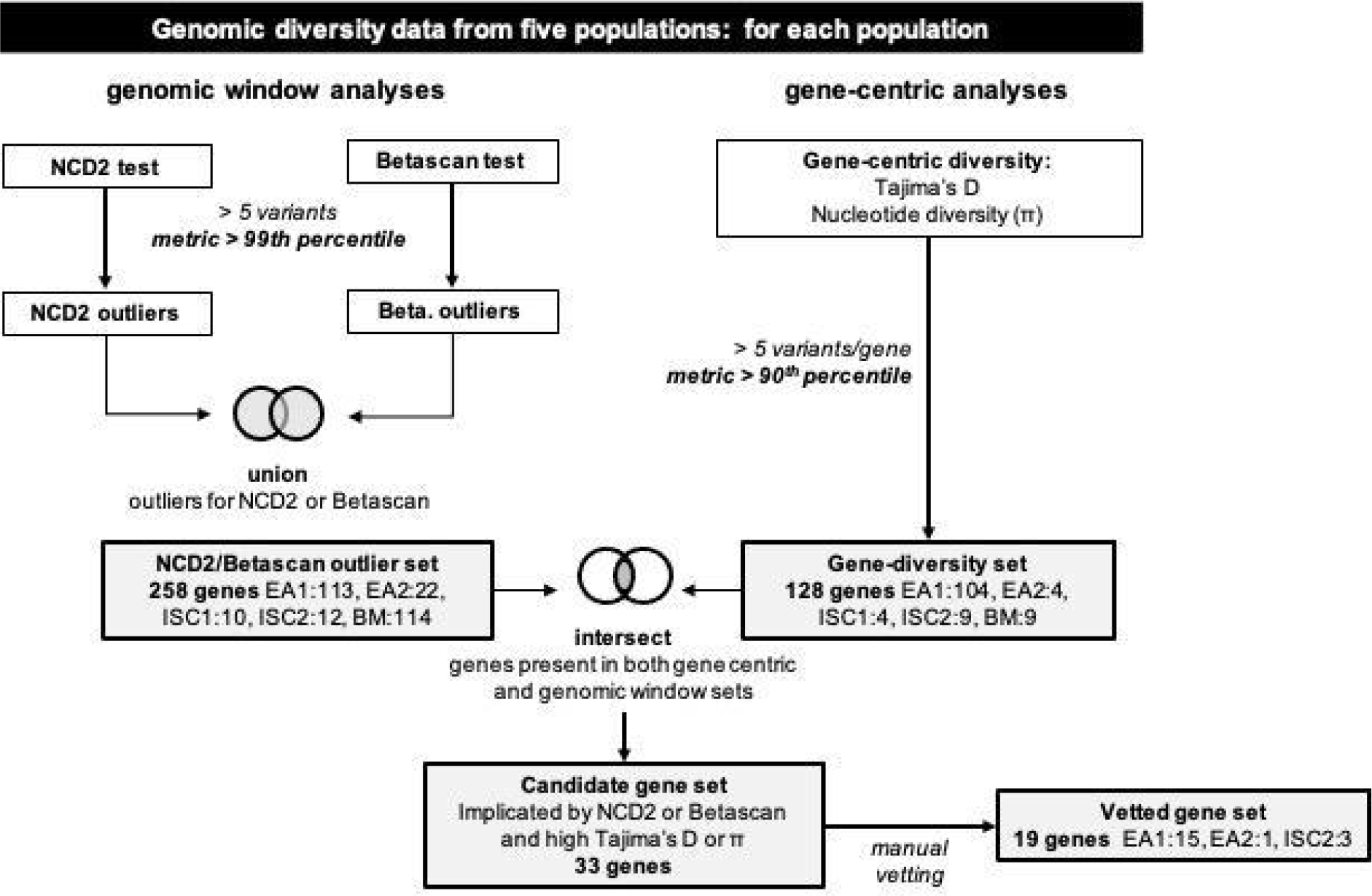
Search strategy employed in balancing selection search. Full description in Supplementary Text 1.

**Supplementary Figure 7.**
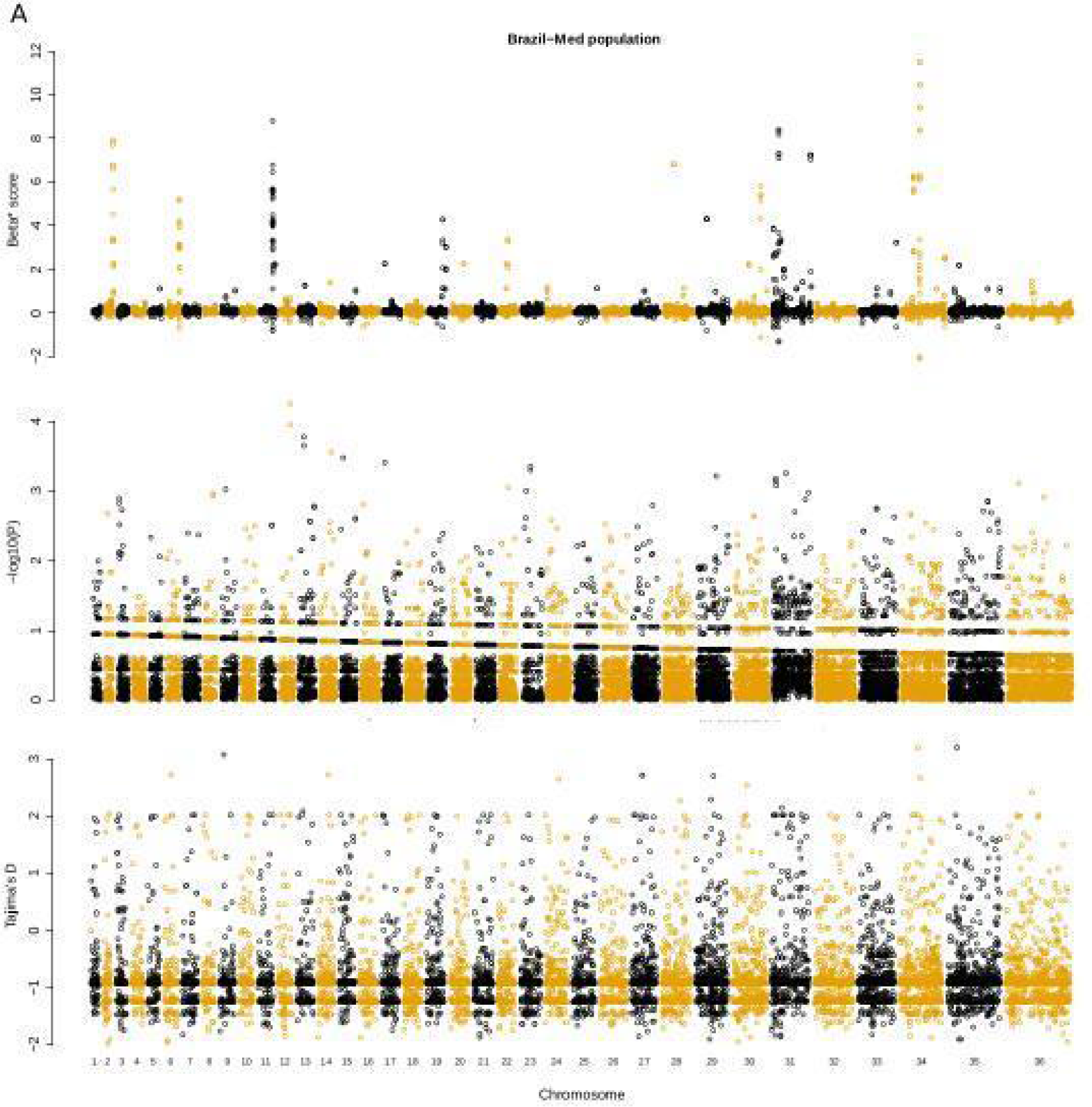
Comparison of balancing selection tests for population BM. Panels from top: Betascan2, NCD2, Tajima’s *D*.

**Supplementary Figure 8.**
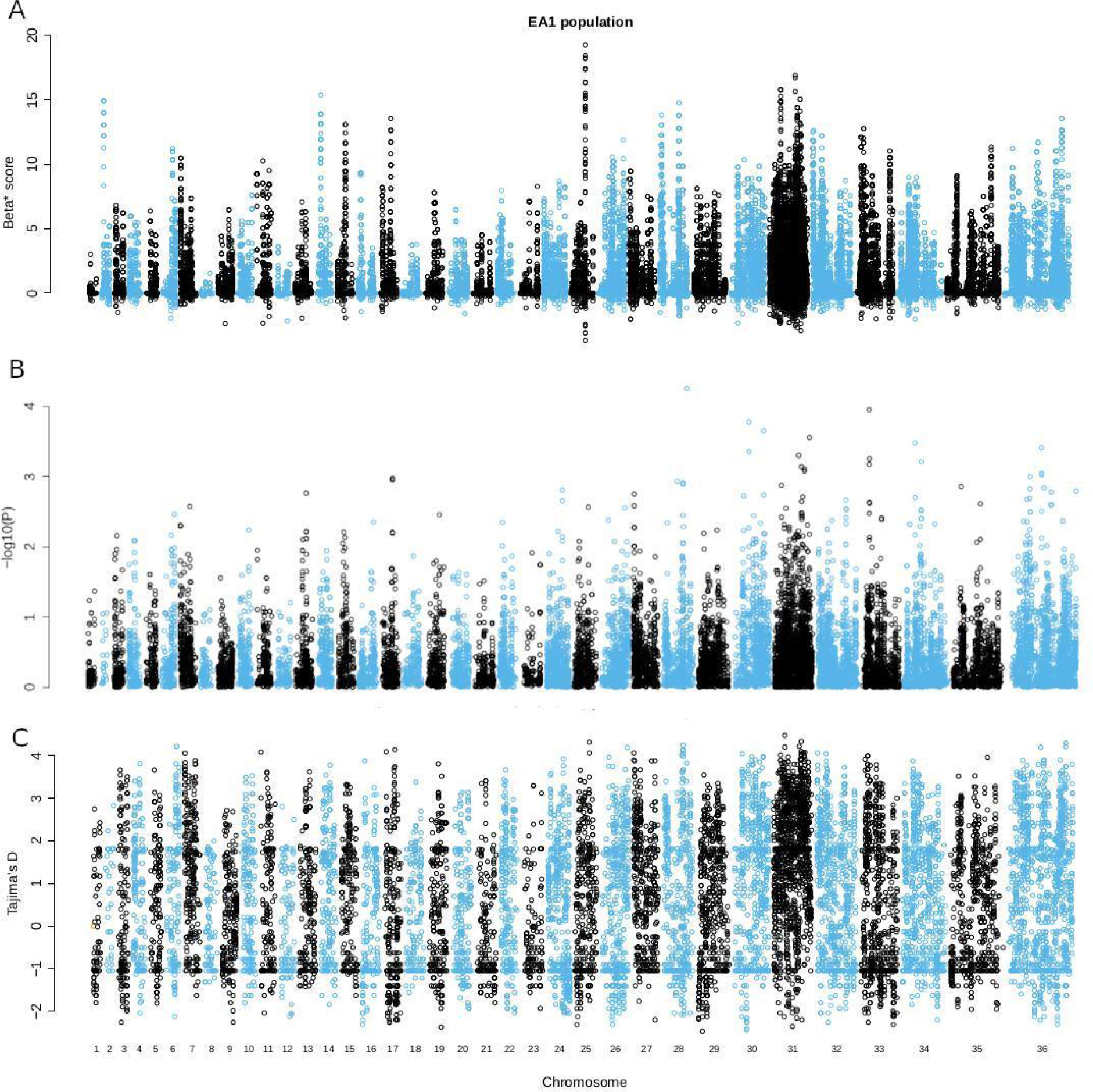
Comparison of balancing selection tests for population EA1. Panels from top: Betascan2, NCD2, Tajima’s *D*.

**Supplementary Figure 9.**
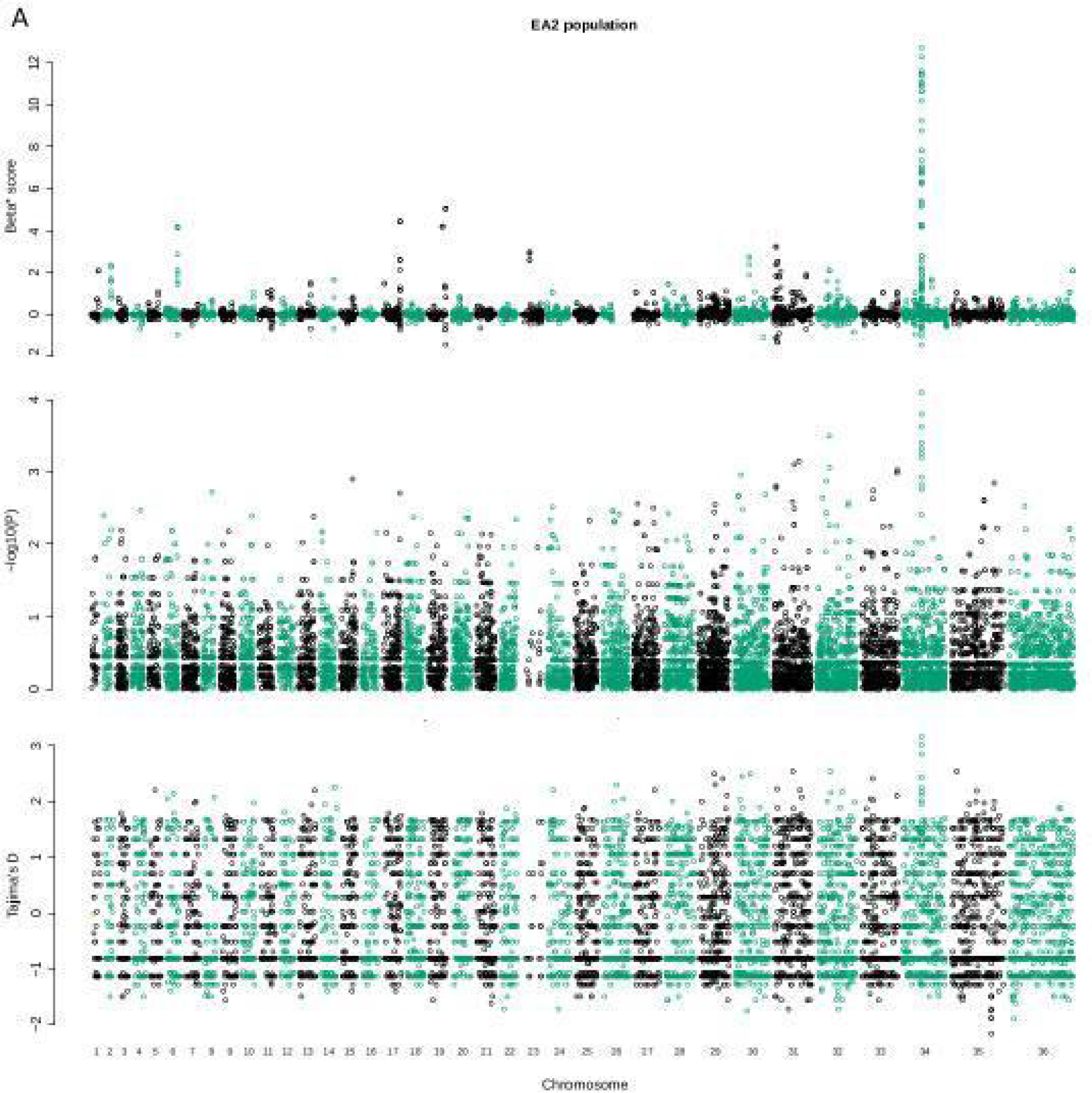
Comparison of balancing selection tests for population EA2. Panels from top: Betascan2, NCD2, Tajima’s *D*.

**Supplementary Figure 10.**
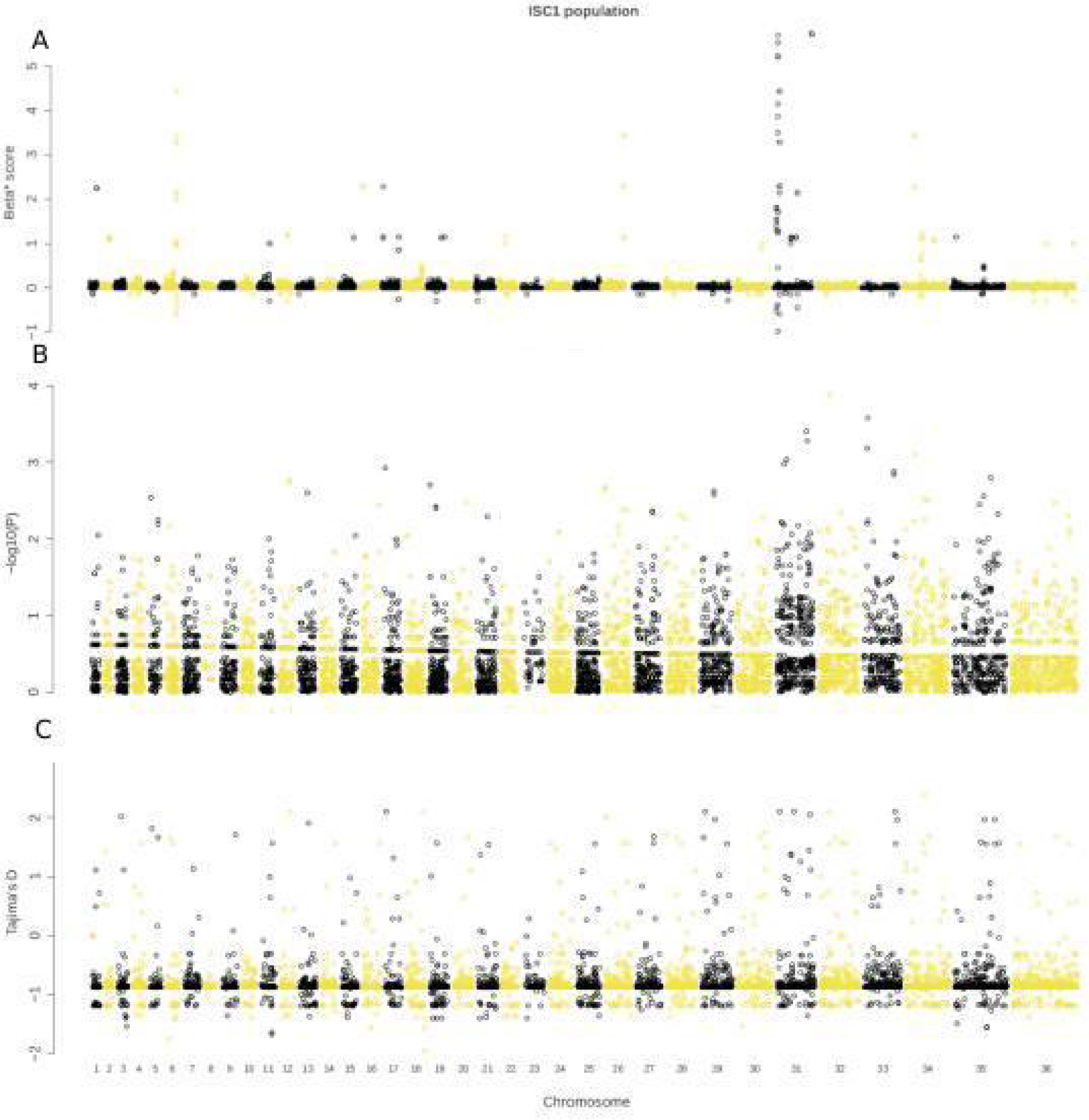
Comparison of balancing selection tests for population ISC1. Panels from top: Betascan2, NCD2, Tajima’s *D*.

**Supplementary Figure 11.**
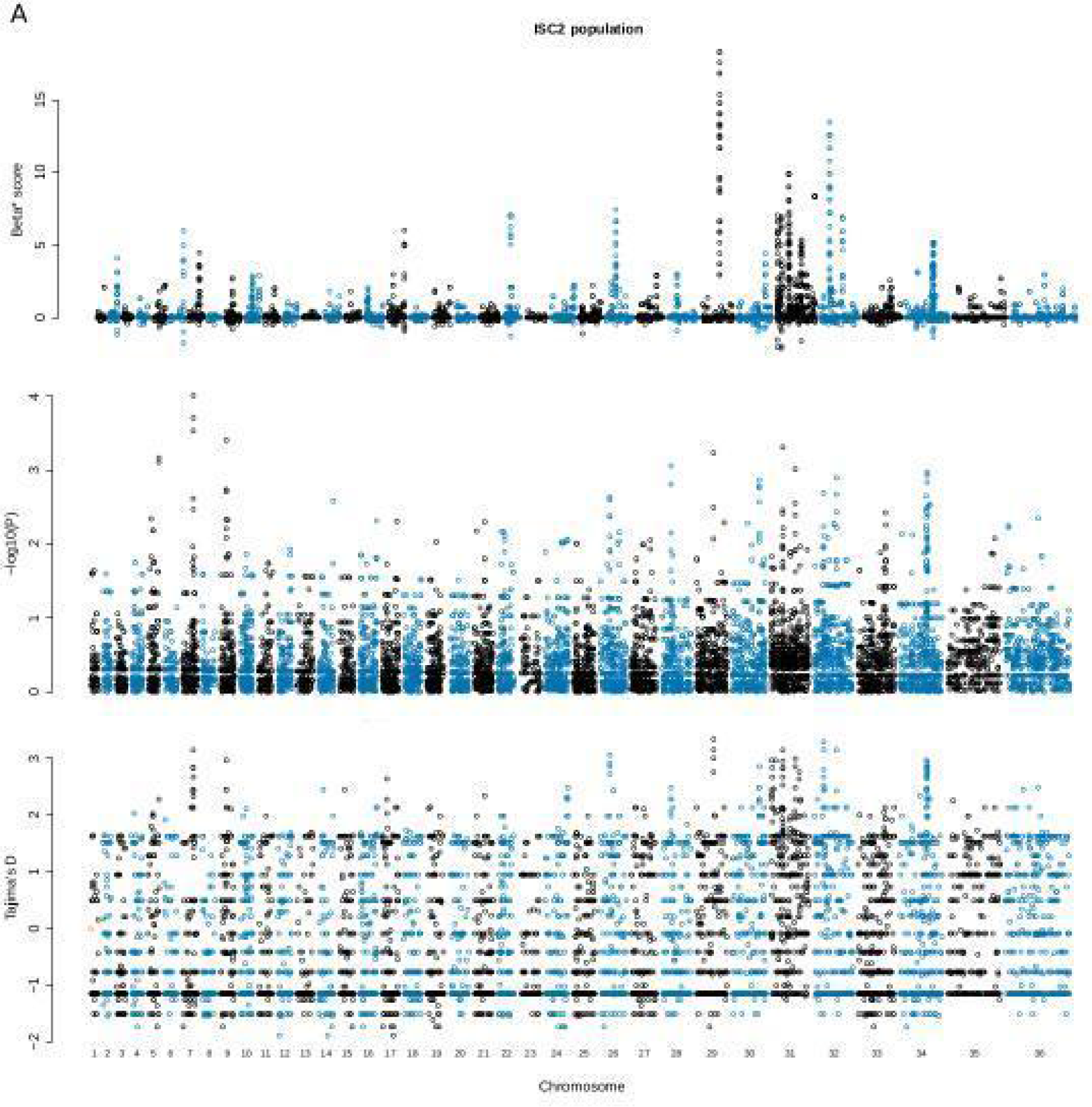
Comparison of balancing selection tests for population ISC2. Panels from top: Betascan2, NCD2, Tajima’s *D*.

**Supplementary Figure 12.**
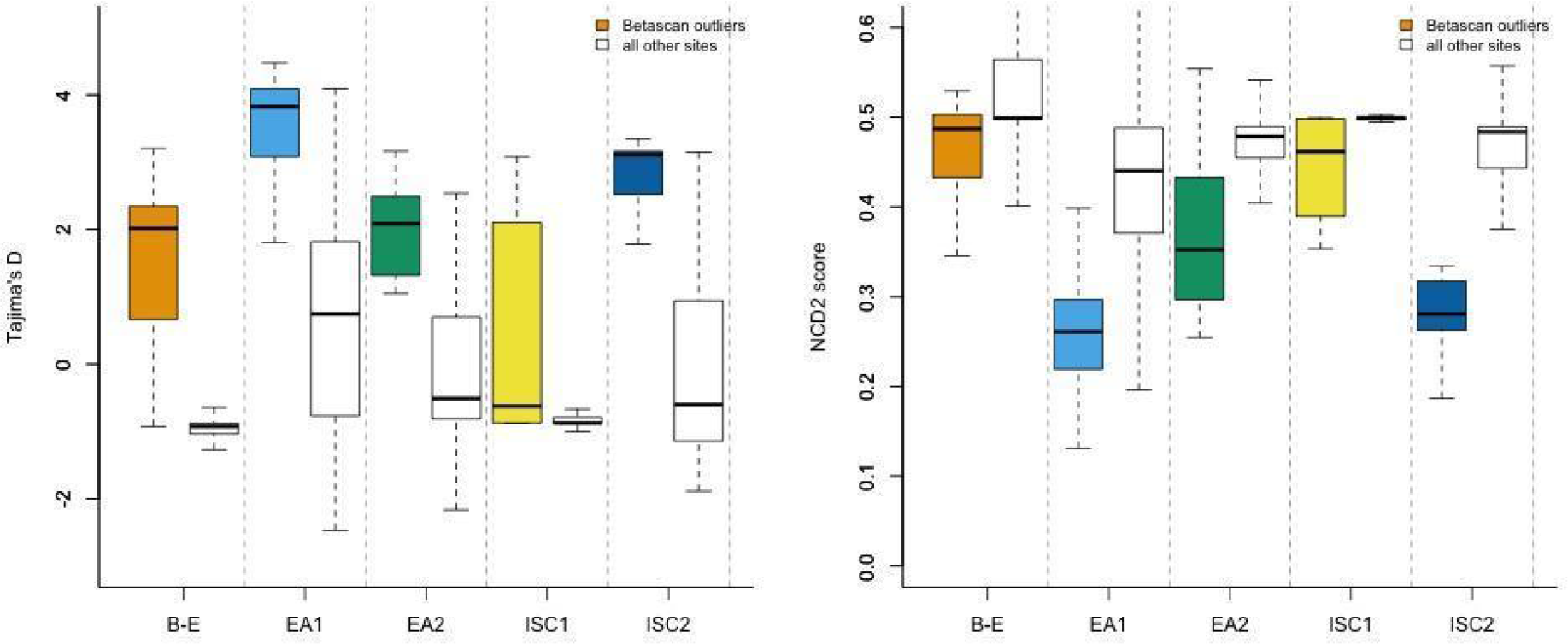
Consistency between tests for balancing selection. In each population, *Betascan* outliers in the 99th percentile of the *β*^(2)^ statistic (Siewert and Voight 2020) were enriched for high Tajima’s *D* (left panel) values, and low *NCD2* scores (right panel). Note that *NCD2* scores are expected to be *low* for sites subject to BS (Bitarello et al. 2018), while Tajima’s *D* will return *higher* values for BS sites.

**Supplementary Figure 13.**
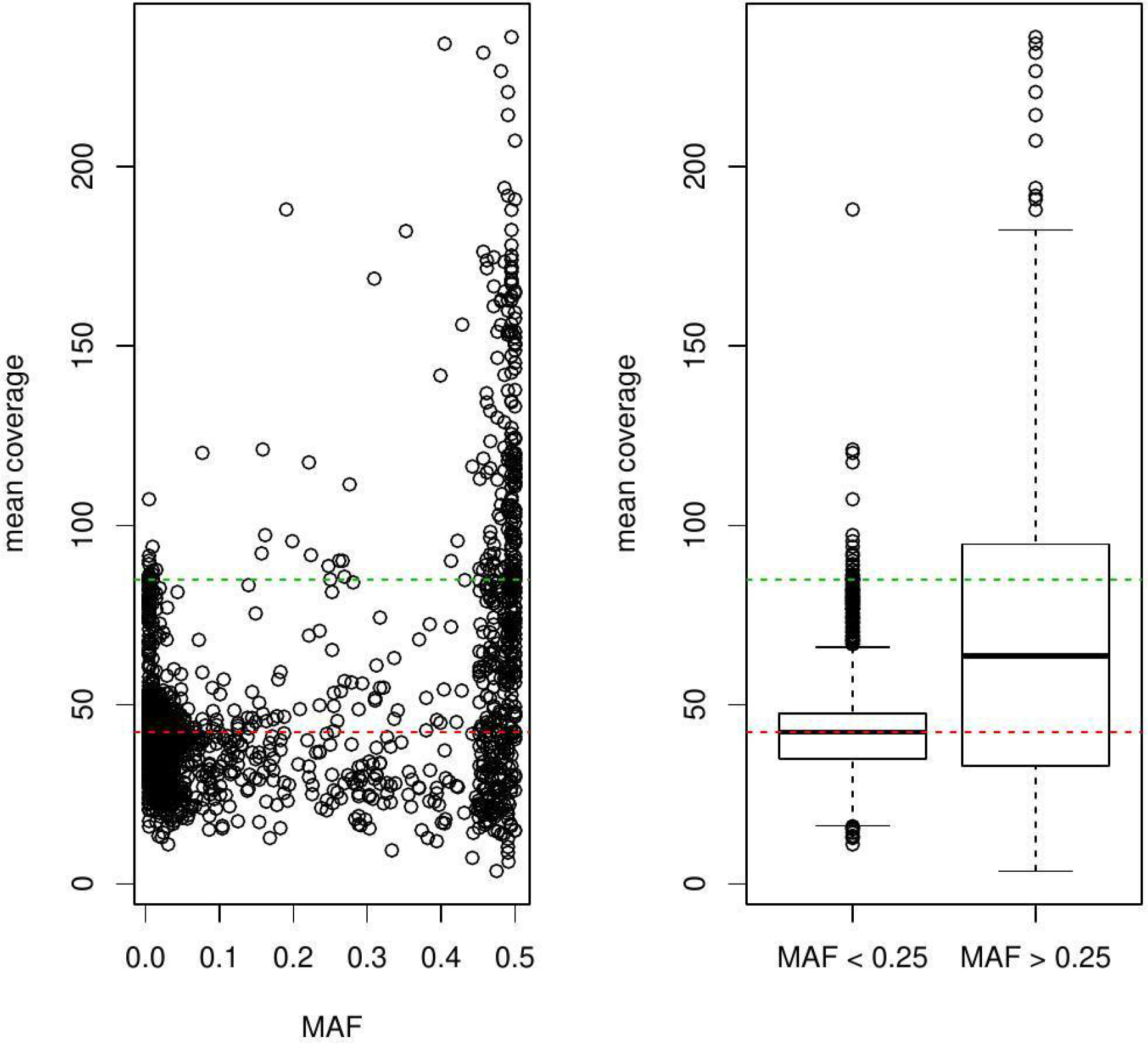
Correlation between MAF and read coverage at SNP sites. Modelling showed that duplications resulted in a systematic bias against calling rare alleles. We removed any SNP/indel sites where the mean variant coverage within the ADMIXTURE-defined population was ≥1.5x larger than the median coverage (corresponding to triploid sites in a generally diploid chromosome), or ≥1.25x larger than the median coverage for chromosome 31 (corresponding to tetraploid sites in a generally triploid chromosome). We also removed sites where coverage was highly variable, by excluding sites in the upper 5th percentile of the standard deviation.

**Supplementary Figure 14.**
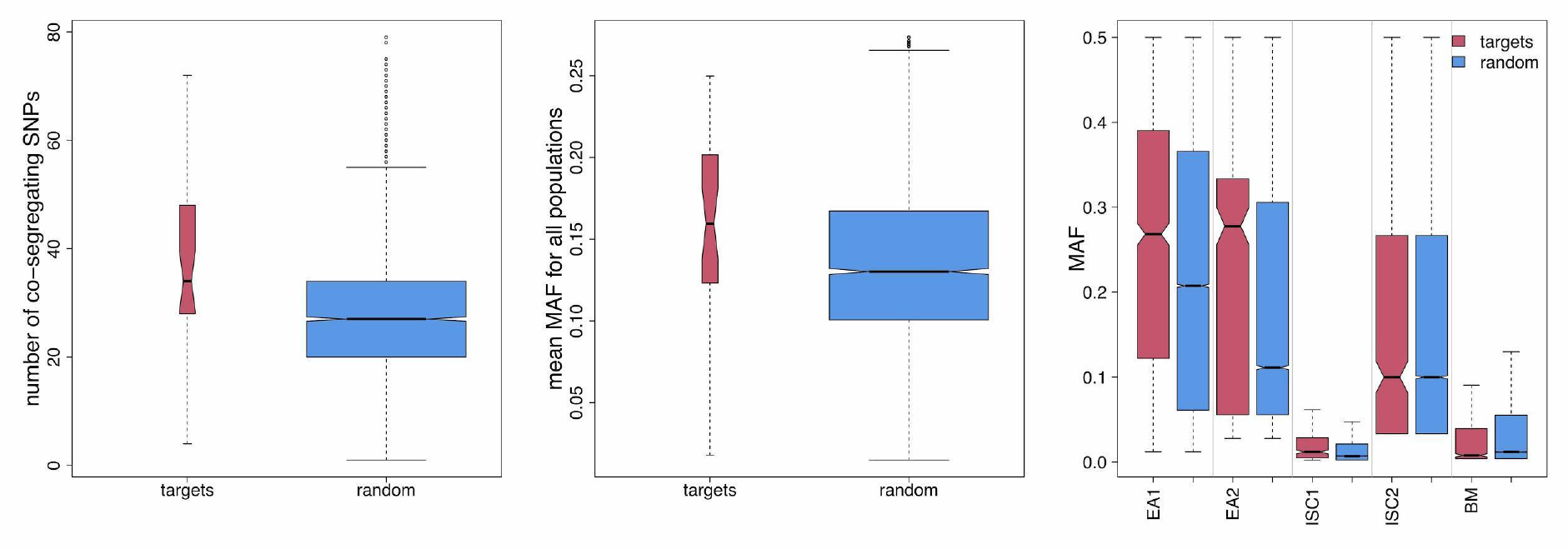
Candidates for BS are enriched for high MAF co-segregating SNPs. Left panel; for the 33 candidate BS genes, we show the number of variants (SNPs/indels) in 130 kb proximal genomic windows that segregate between more than one population, compared to 330 length- and chromosome-matched genomic windows chosen at random. These distributions are significantly different (Mann-Whitney *P* = 0.0001118). Middle panel, these variants have higher mean minor allele frequencies (MAF, means calculated between all populations) compared to matched random controls (Mann-Whitney *P* = 0.001374). Right panel, comparison of the MAF of co-segregating SNPs between populations indicates that target MAF is significantly elevated in the EAst African populations EA1 (where most targets were discovered) and EA2.

**Supplementary Figure 15.**
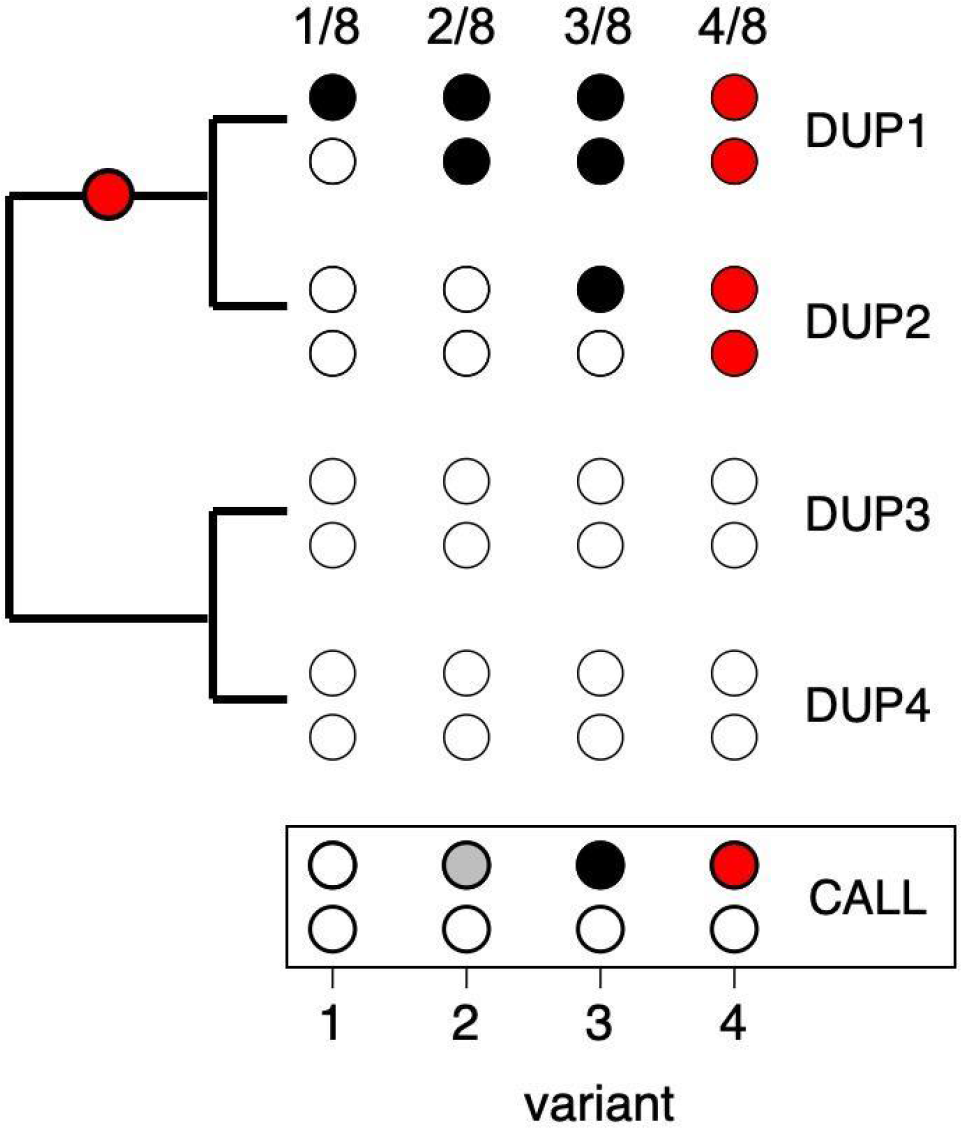
Systematic under-calling of rare alleles in duplicated genes. We consider variant calling in one strain, in a gene that contains four copies, and four variant positions. Duplications have arisen with some arbitrary phylogeny. In a diploid chromosome, there are eight haplotypes. The likely consensus variant call by a variant-caller that assumes a single diploid site is shown in the box below. *Loss of rare alleles*. Rare alleles, such as singletons that have arisen in one copy (variant 1) will be represented by ⅛th of the reads, so are unlikely to be called (two open dots in consensus call). More common alleles, such as those homozygous in one duplication (variant 2), may or not be called, depending on details of variant calling, and stochastic read counts for the alleles (grey dot in consensus call). More common variants in duplications are increasingly likely to be called as heterozygous sites in the consensus call (variant 3). *Appearance of balanced alleles*. In the rare case that a variant becomes fixed in one clade of the duplication phylogeny (variant 4, and red dot in phylogeny), it will be called as a heterozygous site. Depending on the age of the duplication, such a site may be called as a heterozygous site in many/all strains. This will produce the appearance of a balanced polymorphism.

**Supplementary Figure 16. Vignettes of all candidate genes.** Panels from top: Betascan2, NCD2, Tajima’s *D*, nucleotide diversity, MAF and coverage. Note to reviewers: this figure has been uploaded to figshare.

**Supplementary Figure 17.**
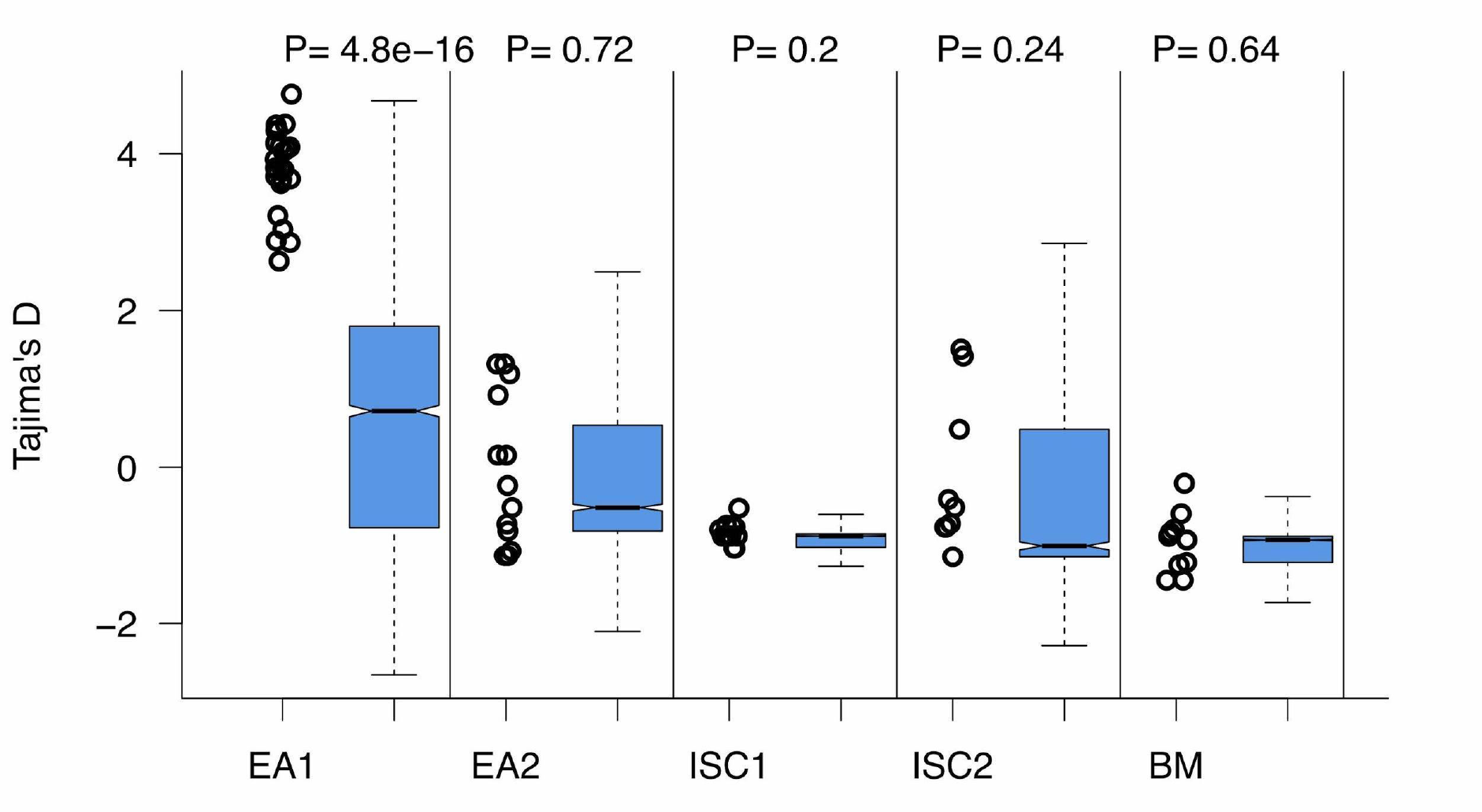
For the 24 candidate BS genes that were discovered in the EA1 population, we show the Tajima *D* values in all populations (open circles), relative to the genome-wide distribution (blue boxes). All p-values are Mann-Whitney tests of candidate BS genes compared to all other genes.

**Supplementary Figure 18.**
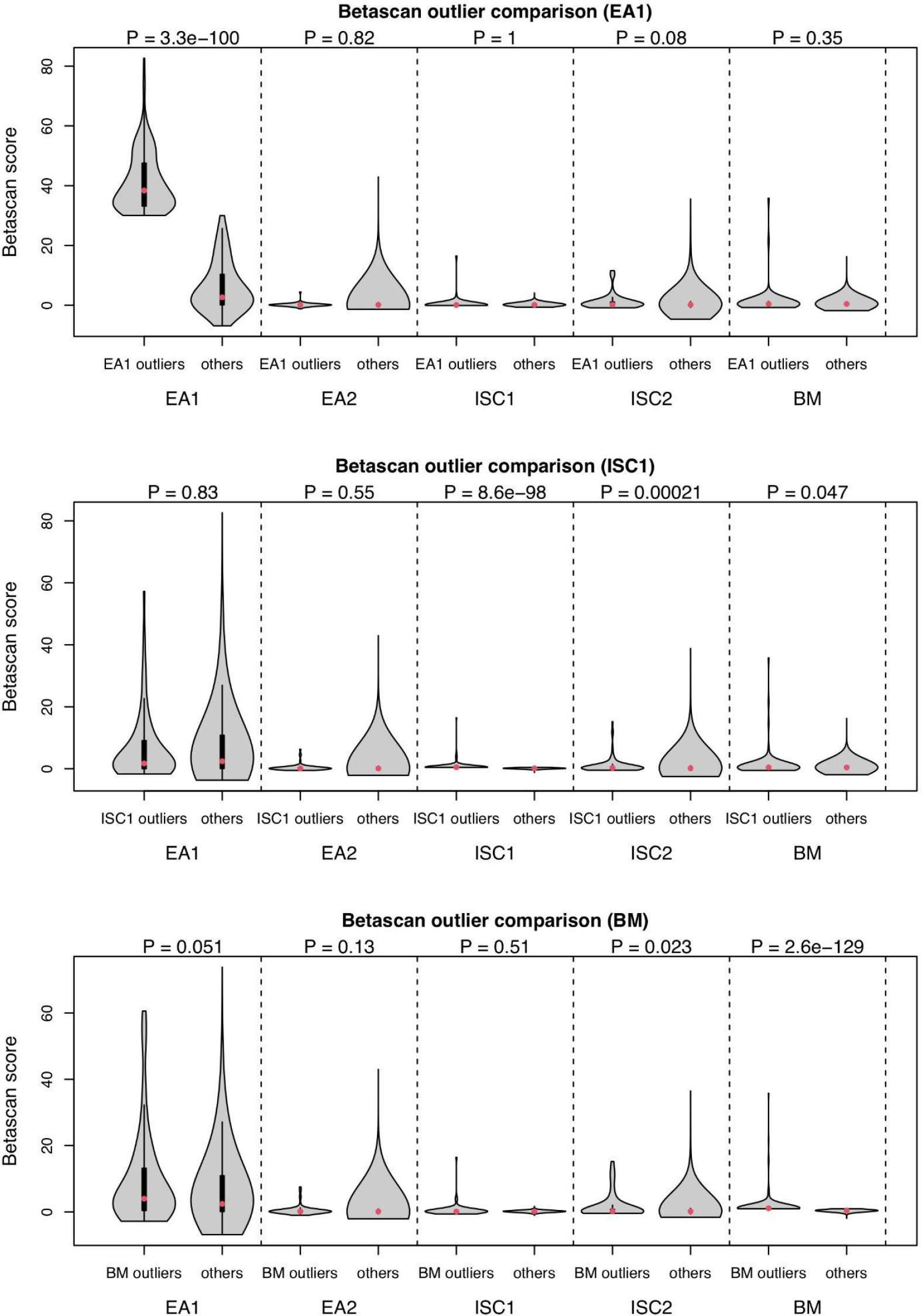
Detecting signals of weak polygenic BS that are consistent between populations. For each of three populations (EA1, ISC1, BM), we identify the 5% outlier genes for the *Betascan2* metric. We then examine whether these genes have significantly higher *Betascan2* scores in any other population (e.g. are *Betascan2* outliers from EA1 outliers in any other population). All comparisons use Wilcoxon signed rank tests to compare outlier genes to all other genes within the same population. Upper panel: *Betascan2* outliers from EA1 do not have higher *Betascan2* scores in any other population. Middle panel: outliers in ISC1 have elevated scores in ISC2 (P =2 x 10^-4^) and marginally-significant elevation of scores in BM (P = 0.047). Lower panel: outliers in BM show marginally-significant elevation of scores in EA1 and ISC2 (P = 0.05, P = 0.02). Only the ISC1-ISC2 enrichment passes a Bonferroni-corrected P-value threshold of 0.0125 (0.05/4).

## Supplementary Tables

**Supplementary Table 1.** List of isolates (note to reviewers: isolates sequenced in this study have temporary sample accession numbers assigned by NCBI. We will update this before publication and when data have been released).

**Supplementary Table 2.** FST values for each population.

**Supplementary Table 3.** Genes that are *Betscan2* outliers in multiple populations.

**Supplementary Table 4.** Candidate gene set and justifications.

**Supplementary Table 5.** Nucleotide diversity and Tajima’s *D* values for all genes in all populations.

**Supplementary Table 6.** Annotated variants in candidate genes.

